# An open dataset of *Plasmodium falciparum* genome variation in 7,000 worldwide samples

**DOI:** 10.1101/824730

**Authors:** Richard D Pearson, Roberto Amato, Dominic P Kwiatkowski, MalariaGEN *Plasmodium falciparum* Community Project

## Abstract

MalariaGEN is a data-sharing network that enables groups around the world to work together on the genomic epidemiology of malaria. Here we describe a new release of curated genome variation data on 7,000 *Plasmodium falciparum* samples from MalariaGEN partner studies in 28 malaria-endemic countries. High-quality genotype calls on 3 million single nucleotide polymorphisms (SNPs) and short indels were produced using a standardised analysis pipeline. Copy number variants associated with drug resistance and structural variants that cause failure of rapid diagnostic tests were also analysed. Almost all samples showed genetic evidence of resistance to at least one antimalarial drug, and some samples from Southeast Asia carried markers of resistance to six commonly-used drugs. Genes expressed during the mosquito stage of the parasite life-cycle are prominent among loci that show strong geographic differentiation. By continuing to enlarge this open data resource we aim to facilitate research into the evolutionary processes affecting malaria control and to accelerate development of the surveillance toolkit required for malaria elimination.

## Introduction

A major obstacle to malaria elimination is the great capacity of the parasite and vector populations to evolve in response to malaria control interventions. The widespread use of chloroquine and DDT in the 1950’s led to high levels of drug and insecticide resistance, and the same pattern has been repeated for other first-line antimalarial drugs and insecticides. Over the past 15 years, mass distribution of pyrethroid-treated bednets in Africa and worldwide use of artemisinin combination therapy (ACT) has led to substantial reductions in malaria prevalence and mortality, but there are rapidly increasing levels of resistance to ACT in Southeast Asian parasites and of pyrethroid resistance in African mosquitoes. A deep understanding of local patterns of resistance and the continually changing nature of the local parasite and vector populations is necessary to manage the use of drugs and insecticides and to deploy public health resources for maximum sustainability and impact.

Current methods for genetic surveillance of the parasite population are largely based on targeted genotyping of specific loci, e.g. known markers of drug resistance. Whole genome sequencing of malaria parasites is currently more expensive and complex, particularly at the stage of data analysis, but it is an important adjunct to targeted genotyping, as it provides a more comprehensive picture of parasite genetic variation. It is particularly important for discovery of new drug resistance markers and for monitoring patterns of gene flow and evolutionary adaptation in the parasite population.

The *Plasmodium falciparum* Community Project (*Pf* Community Project) was established with the aim of integrating parasite genome sequencing into clinical and epidemiological studies of malaria (www.malariagen.net/projects). It forms part of the Malaria Genomic Epidemiology Network (MalariaGEN), a global data-sharing network comprising multiple partner studies, each with its own research objectives and led by a local investigator.^1^ Genome sequencing was performed centrally, and partner studies were free to analyse and publish the genetic data produced on their own samples, in line with MalariaGEN’s guiding principles on equitable data sharing.^1–3^ A programme of capacity building for research into parasite genetics was developed at multiple sites in Africa alongside the *Pf* Community Project.^4^

The first phase of the project focused on developing simple methods to obtain purified parasite genome DNA from small blood samples collected in the field ^5,6^ and on establishing reliable computational methods for variant discovery and genotype calling from short-read sequencing data.^7^ This presented a number of analytical challenges due to long tracts of highly repetitive sequence and hypervariable regions within the *P. falciparum* genome, and also because a single infection can contain a complex mixture of genotypes. Once a reliable analysis pipeline was in place, a process was established for periodic data releases to partners, with continual improvements in data quality as new analytical methods were developed.

Data from the *Pf* Community Project were initially released through a companion project called Pf3k (www.malariagen.net/data), whose goal was to bring together leading analysts from multiple institutions to benchmark and standardise methods of variant discovery and genotyping calling. A visual analytics web application was developed^8^ for researchers to explore the data (www.malariagen.net/apps/pf3k). The open dataset was enlarged in 2016 when multiple partner studies contributed to a consortial publication on 3,488 samples from 23 countries.^9^

Data produced by the *Pf* Community Project have been used to address a broad range of research questions, both by the groups that generated samples and data and by the wider research community, and have generated over 50 previous publications (refs 5-55). These data have become a key resource for the epidemiology and population genetics of antimalarial drug resistance^9–22^ and an important platform for the discovery of new genetic markers and mechanisms of resistance through genome-wide association studies^23–27^ and combined genome-transcriptome analysis. ^28^ The data have also been used to study gene deletions that cause failure of rapid diagnostic tests^29^; to characterise genetic variation in malaria vaccine antigens ^30,31^; to screen for new vaccine candidates^32^ ; to investigate specific host-parasite interactions^33,34^; and to describe the evolutionary adaptation and diversification of local parasite populations. ^7,9,12,35–40^

The *Pf* Community Project data also provide an important resource for developing and testing new analytical and computational methods. A key area of methods development is quantification of within-host diversity ^7,41–46^, estimation of inbreeding^7,47^, and deconvolution of mixed infections into individual strains.^48,49^ The data have also been used to develop and test methods for estimating identity by descent^50,51^, imputation^52^, typing structural variants^53^, designing other SNP genotyping platforms^54^ and data visualisation^8,55^. In a companion study we performed whole genome sequencing of experimental genetic crosses of *P. falciparum*, and this provided a benchmark to test the accuracy of our genotyping methods, and to conduct an in-depth analysis of indels, structural variants and recombination events which are complicated to ascertain in these population genetic samples^56^.

Here we describe a new release of curated genome variation data on 7,113 samples of *P. falciparum* collected by 49 partner studies from 73 locations in Africa, Asia, South America and Oceania between 2002 and 2015 (Table 1, Supplementary Table 1 and 2).

**Table 1.** Count of samples in the dataset. Countries are grouped into eight geographic regions based on their geographic and genetic characteristics. For each country, the table reports: the number of distinct sampling locations; the total number of samples sequenced; the number of high-quality samples included in the analysis; and the percentage of samples collected between 2012-2015, the most recent sampling period in the dataset. Eight samples were obtained from travellers returning from an endemic country, but where the precise site of the infection could not be determined. These were reported from Ghana (3 sequenced samples/2 analysis set samples), Kenya (2/1), Uganda (2/1) and Mozambique (1/1). “Lab samples” contains all sequences obtained from long-term *in vitro* cultured and adapted isolates, e.g. laboratory strains. The breakdown by site is reported in Supplementary table 1 and the list of contributing studies in Supplementary table 2.

| Region | Country | Sampling locations | Sequenced samples | Analysis set samples | % analysis samples 2012-2015 |
| --- | --- | --- | --- | --- | --- |
| **South America (SAM)** | **Colombia** | 4 | 16 | 16 | 0% |
|  | **Peru** | 2 | 23 | 21 | 0% |
| **West Africa (WAF)** | **Benin** | 1 | 102 | 36 | 100% |
|  | **Burkina Faso** | 1 | 57 | 56 | 0% |
|  | **Cameroon** | 1 | 239 | 235 | 100% |
|  | **Gambia** | 4 | 277 | 219 | 67% |
|  | **Ghana** | 3 | 1,003 | 849 | 56% |
|  | **Guinea** | 2 | 197 | 149 | 0% |
|  | **Ivory Coast** | 3 | 70 | 70 | 100% |
|  | **Mali** | 5 | 449 | 426 | 80% |
|  | **Mauritania** | 4 | 86 | 76 | 100% |
|  | **Nigeria** | 2 | 42 | 29 | 97% |
|  | **Senegal** | 1 | 86 | 84 | 100% |
| **Central Africa(CAF)** | **Congo DR** | 1 | 366 | 344 | 100% |
| **East Africa (EAF)** | **Ethiopia** | 2 | 34 | 21 | 100% |
|  | **Kenya** | 3 | 129 | 109 | 55% |
|  | **Madagascar** | 3 | 25 | 24 | 100% |
|  | **Malawi** | 2 | 351 | 254 | 0% |
|  | **Tanzania** | 5 | 350 | 316 | 85% |
|  | **Uganda** | 1 | 14 | 12 | 0% |
| **South Asia (SAS)** | **Bangladesh** | 2 | 93 | 77 | 64% |
| **Western Southeast Asia (WSEA)** | **Myanmar** | 5 | 250 | 211 | 71% |
|  | **Western Thailand** | 2 | 962 | 868 | 24% |
| **Eastern Southeast Asia (ESEA)** | **Cambodia** | 5 | 1,214 | 896 | 32% |
|  | **Northeastern Thailand** | 1 | 28 | 20 | 75% |
|  | **Laos** | 2 | 131 | 120 | 21% |
|  | **Viet Nam** | 2 | 264 | 226 | 11% |
| **Oceania (OCE)** | **Indonesia** | 1 | 92 | 80 | 73% |
|  | **Papua New Guinea** | 3 | 139 | 121 | 63% |
| **Returning travellers** | **Various locations** | 0 | 8 | 5 | 0% |
| **Lab samples** | **Various locations** | 0 | 16 | 0 | 0% |
| **Total** |  | **73** | **7,113** | **5,970** | **52%** |

## Results

### Variant discovery and genotyping

We used the Illumina platform to produce genome sequencing data on all samples and we mapped the sequence reads against the *P. falciparum* 3D7 v3 reference genome. The median depth of coverage was 73 sequence reads averaged across the whole genome and across all samples. We constructed an analysis pipeline for variant discovery and genotyping, including stringent quality control filters that took into account the unusual features of the *P. falciparum* genome, incorporating lessons learnt from our previous work^7,56^ and the Pf3k project, as outlined in the Methods section.

In the first stage of analysis we discovered variation at over six million positions, corresponding to about a quarter of the 23Mb *P. falciparum* genome (Supplementary Table 3). These included 3,168,721 single nucleotide polymorphisms (SNPs): these were slightly more common in coding than non-coding regions and were mostly biallelic. The remaining 2,882,975 variants were predominantly short indels but also included more complex combinations of SNPs and indels: these were much more abundant in non-coding than coding regions, and mostly had at least 3 alleles. The predominance of indels in non-coding regions has been previously observed and is most likely a consequence of the extreme AT bias which leads to many short repetitive sequences. ^56,57^

For the purpose of this analysis, we excluded all variants in subtelomeric and internal hypervariable regions, mitochondrial and apicoplast genomes, and some other regions of the genome where the mapping of short sequence reads is prone to a high error rate due to extremely high rates of variation. ^56^ A total of 1,838,733 SNPs (of which 1,626,886 were biallelic) and 1,276,027 indels (or SNP/indel combinations) passed all these filters. The pass rate for SNPs in coding regions (66%) was considerably higher than that for SNPs in non-coding regions (47%), indels in coding regions (37%) and indels in non-coding regions (47%). Finally we removed samples with a low genotyping success rate or other quality control issues. We also removed replicates and 41 samples with genetic markers of infection by multiple *Plasmodium* species, leaving 5,970 high-quality samples from 28 countries (Table 1).

We used coverage and read pair analysis to determine duplication genotypes around *mdr1*, *plasmepsin2/3* and *gch1*, each of which are associated with drug resistance. For each of these three genes we discovered many different sets of breakpoints (29, 10 and 3 pairs of breakpoints for *mdr1*, *gch1*, and *plasmepsin 2/3*, respectively), including complex rearrangements^58^ that to the best of our knowledge have not been observed before in *Plasmodium* species (Supplementary Note, Supplementary Tables 4-6). We also used sequence reads coverage to identify large structural variants that appear to delete or disrupt *hrp2* and *hrp3*, an event that can cause rapid diagnostic tests to malfunction.

The population genetic analyses in this paper are based on the filtered dataset of high-quality SNP genotypes in 5,970 samples. These data are openly available, together with annotated genotyping data on 6 million putative variants in all 7,113 samples, plus details of partner studies and sampling locations, at www.malariagen.net/resource/26 .

### Global population structure

The genetic structure of the global parasite population reflects its geographic regional structure ^7,9,10^ as illustrated by a neighbour-joining tree and a principal component analysis of all samples based on their SNP genotypes (Figure 1). Based on these observations we grouped the samples into eight geographic regions: West Africa, Central Africa, East Africa, South Asia, the western part of Southeast Asia, the eastern part of Southeast Asia, Oceania and South America. Each of these can be viewed as a regional sub-population of parasites, which is more or less differentiated from other regional sub-populations depending on rates of gene flow and other factors. The different regions encompass a range of epidemiological and environmental settings, varying in transmission intensity, vector species and history of antimalarial drug usage. Note these regional classifications are intentionally broad, and therefore overlook many interesting aspects of local population structure, e.g. a distinctive Ethiopian sub-population can be identified by more detailed analysis of African samples.^12^

**Figure 1.**
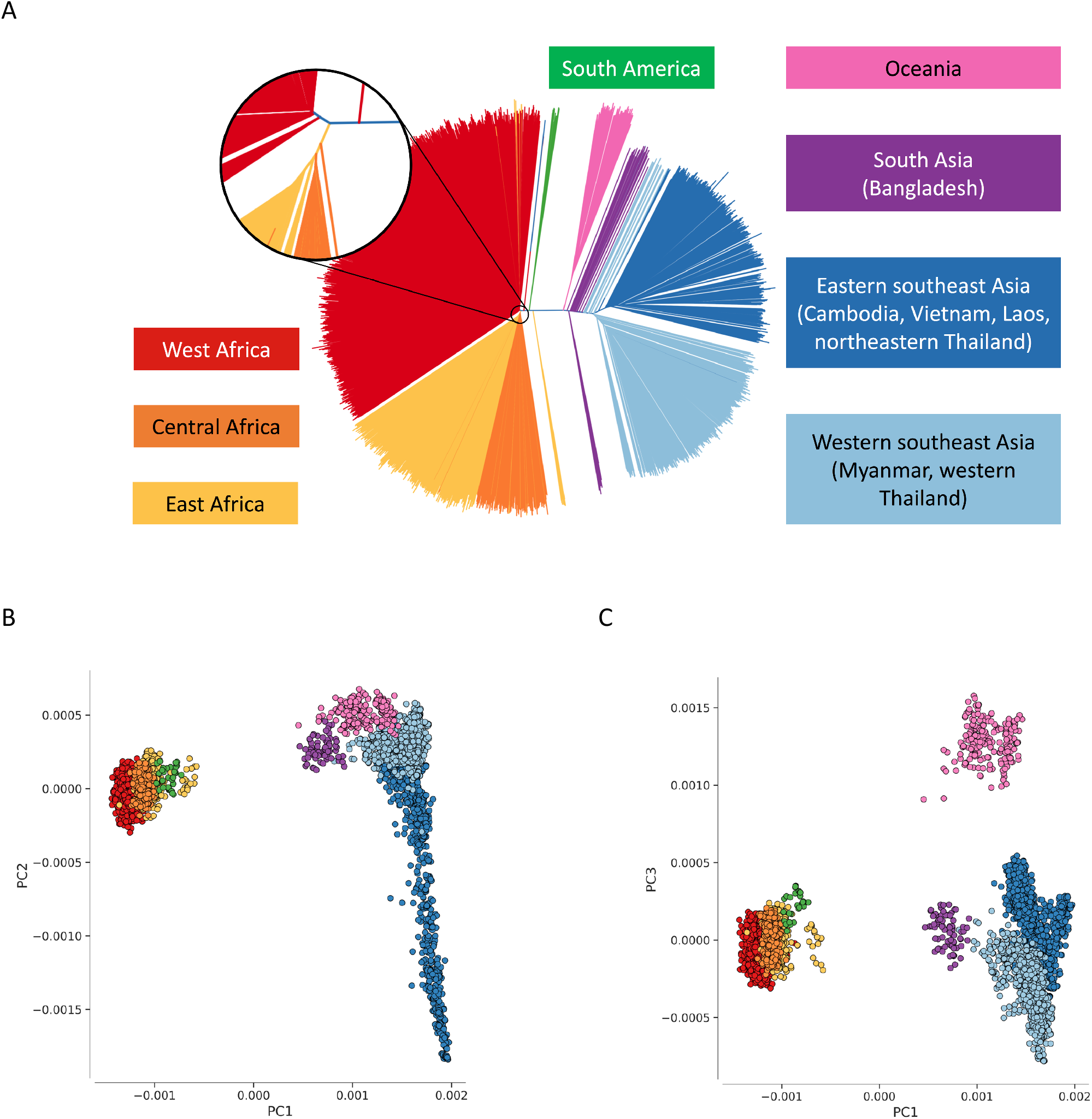
Population structure. (A) Genome-wide unrooted neighbour-joining tree showing population structure across all sites, with sample branches coloured according to country groupings (Table 1): South America (green, n=37); West Africa (red, n=2231); Central Africa (orange, n=344); East Africa (yellow, n=739); South Asia (purple, n=77); West south-east Asia (light blue; n=1079); East south-east Asia (dark blue; n=1262); Oceania (magenta; n=201). The circular inset shows a magnified view of the part of the tree where the majority of samples from Africa coalesce, showing that the three African sub-regions are genetically close but distinct. (B, C) First three component of a genome-wide principal coordinate analysis. The first axis (PC1) captures the separation of African and South American from Asian samples. The following two axes (PC2 and PC3) capture finer levels of population structure due to geographical separation and selective forces. Each point represents a sample and the colour legend is the same as above.

Genetically mixed infections were considerably more common in Africa than other regions, consistent with the high intensity of malaria transmission in Africa (Figure 2a). Analysis of *F*_*WS*_, a measure of within-host diversity^7^, shows that most samples from Southeast Asia (1763/2341), South America (37/37) and Oceania (158/201) have *F*_*WS*_ *>*0.95, which to a first approximation indicates that the infection is dominated by a clonal population of parasite.^41^ In contrast, nearly half of samples from Africa (1625/3314) have *F*_*WS*_ <0.95, indicating the presence of more complex infections. Genetically mixed infections were also common in Bangladesh (41/77 samples have *F*_*WS*_ <0.95), another area of high malaria transmission and the only South Asian country represented in this dataset, but did not reach the extremely high levels of within-host diversity (*F*_*WS*_ <0.2) observed in some samples from Africa.

**Figure 2.**
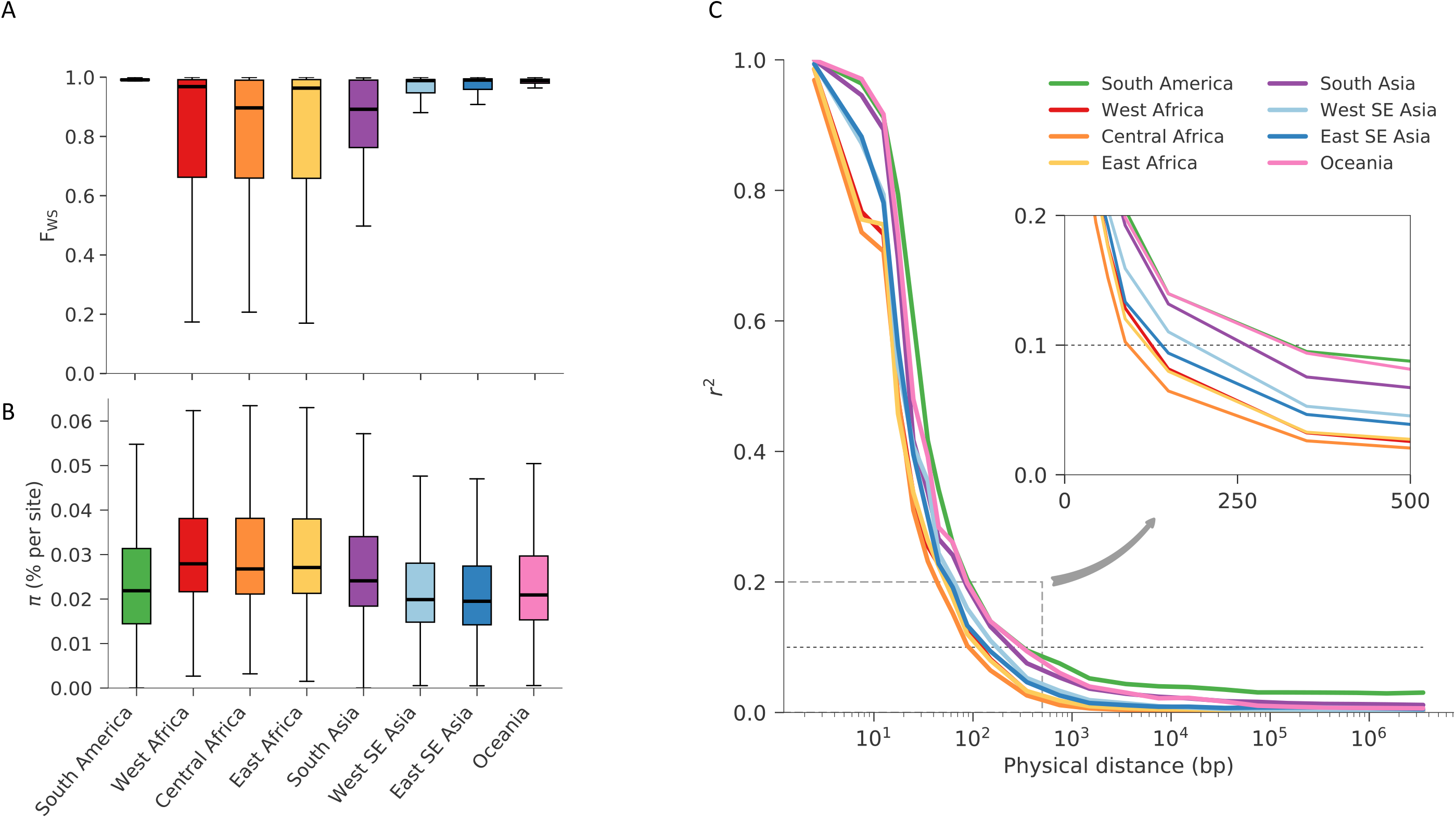
Characteristics of the eight regional parasite populations. (A) Distribution of within-host diversity, as measured by F_WS_, showing that genetically mixed infections were considerably more common in Africa than other regions, consistent with the high intensity of malaria transmission in Africa. (B) Distribution of per site nucleotide diversity calculated in non-overlapping 25kbp genomic windows. We only considered coding biallelic SNPs to reduce the ascertainment bias caused by poor accessibility of non-coding regions. In both previous panels, thick lines represent median values, boxes show the interquartile range, and whiskers represent the bulk of the distribution, discounting outliers. (C) Genome-wide median LD (y-axis, measured by r^2^) between pairs of SNPs as function of their physical distance (x-axis, in bp), showing a rapid decay in all regional parasite populations. The inset panel shows a magnified view of the decay, showing that in all populations r^2^ decayed below 0.1 (dashed horizontal line) within 500 bp. All panels utilise the same palette, with colours denoting each geographic region.

The average nucleotide diversity across the global sample collection was 0.040% (median=0.028%), i.e. two randomly-selected samples differ by an average of 4 nucleotide positions per 10kb. Levels of nucleotide diversity vary greatly across the genome^56^ and also geographically (Figure 2b). Distributions of values were highest in Africa, followed by Bangladesh, but the scale of regional differences was relatively modest, ranging from an average of 0.030% in Eastern Southeast Asia to 0.040% in West Africa (median=0.019% and 0.028% respectively; Figure 2b). In other words, the nucleotide diversity of each regional parasite population was not much less than that of the global parasite population. This is consistent with the idea that the global *P. falciparum* population has a common African origin and that historically there must have been significant levels of migration.

All regional sub-populations showed very low levels of linkage disequilibrium relative to human populations, e.g. r^2^ decayed to <0.1 within 500 bp (Figure 2c). As expected, African populations had the highest rates of LD decay, implying the highest levels of haplotype diversity.

### Geographic patterns of population differentiation and gene flow

Parasite sub-populations in different locations naturally tend to differentiate over time unless there is sufficient gene flow to counterbalance genetic drift. Genome-wide estimates of *F*_*ST*_ provide an indicator of this process of genetic differentiation, which is partly determined by geographic distance (Figure 3). For example, we observe much greater genetic differentiation between South America and South Asia (genome-wide average *F*_*ST*_ 0.22) or between Africa and Oceania (0.20) than between sub-regions within Asia (<0.1) or within Africa (<0.02).

**Figure 3.**
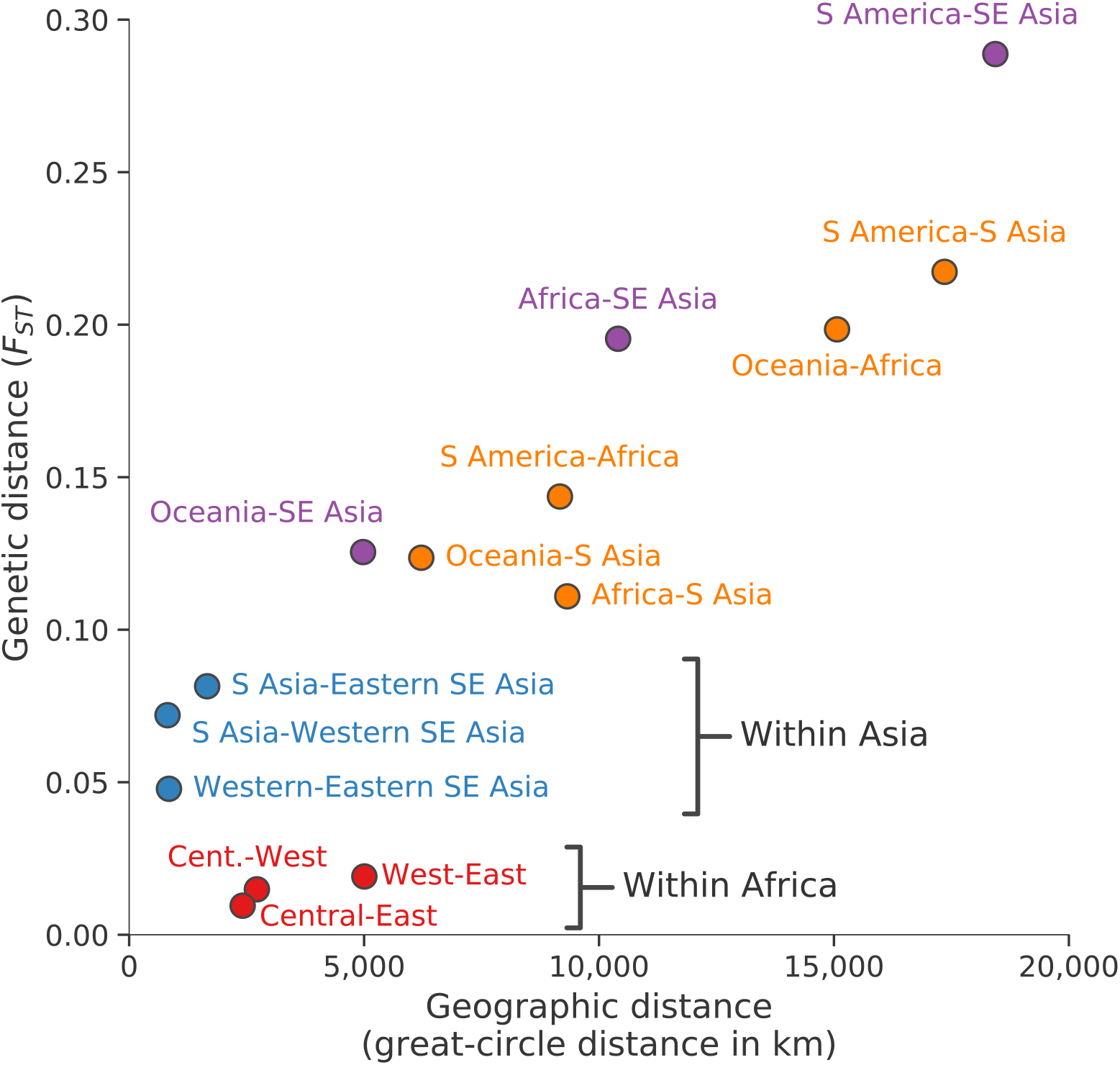
Geographic patterns of population differentiation and gene flow. Each point represents one pairwise comparison between two regional parasite populations. The x-axis reports the geographic separation between the two populations, measured as great-circle distance between the centre of mass of each population and without taking into account natural barriers. The y-axis reports the genetic differentiation between the two populations, measured as average genome-wide F_ST_. Points are coloured based on the regional populations they represent: between African populations (red); between Asian populations (blue); between Southeast Asia (as a whole) and Oceania, Africa or South America (purple); all the rest (orange).

These data reveal some interesting exceptions to the general rule that genome-wide *F*_*ST*_ is correlated with geographic distance. For example, African parasites are more strongly differentiated from Southeast Asian parasites (genome-wide average *F*_*ST*_ 0.20) than they are from parasites in neighbouring Bangladesh (0.11). If this is examined in more detail, there is an unexpectedly steep gradient of genetic differentiation at the geographical boundary between South Asia and Southeast Asia, i.e. parasites sampled in Myanmar and Western Thailand are much more strongly differentiated from parasites sampled in Bangladesh (genome-wide *F*_*ST*_ 0.07) than would be expected given that these are neighbouring countries. As discussed later, Southeast Asia is the global epicentre of antimalarial drug resistance, and these observations add to a growing body of evidence that Southeast Asian parasites have acquired a wide range of genomic features that are likely due to natural selection rather than genetic drift.^23,40^

It is noteworthy that the level of genetic differentiation between western and eastern parts of Southeast Asia (genome-wide *F*_*ST*_ 0.05) is greater than between West Africa and East Africa (0.02) although the geographic distances are much greater in Africa. This is likely due to the lower intensity of malaria transmission in Southeast Asia, and in particular the presence of a malaria-free corridor running through Thailand, which act as barriers to gene flow across the region.^23,40^

### Genes with high levels of geographic differentiation

The *F*_*ST*_ metric can also be calculated for individual variants to identify specific genes that have acquired high levels of geographic differentiation relative to the genome as a whole. This can be done either at the global level (to identify variants that are highly differentiated between different regions of the world) or at the local level (to identify variants that are highly differentiated between different sampling locations within a region).

To identify variants that are strongly differentiated at the global level, we began by estimating *F*_*ST*_ for each SNP across all of the eight regional sub-populations. The group of SNPs with the highest global *F*_*ST*_ levels were found to be strongly enriched for non-synonymous mutations, suggesting that the process of differentiation is at least in part due to natural selection (Figure 4). After ranking all SNPs according to their global *F*_*ST*_ value, we calculated a *global differentiation score* for each gene based on the highest-ranking non-synonymous SNP within the gene (see Methods). All genes are ranked according to their global differentiation score in the accompanying data release, and those with the highest score are listed in Supplementary Table 7. The most highly differentiated gene, *p47*, is known to interact with the mosquito immune system^59^ and has two variants (S242L and V247A) that are at fixation in South America but absent in other geographic regions. Also among the five most highly differentiated genes are *gig* (implicated in gametocytogenesis^60^), *pfs16*, (expressed on the surface of gametes^61^) and *ctrp* (expressed on the ookinete cell surface and essential for mosquito infection^62^). Thus four of the five most highly differentiated parasite genes are involved in the process of transmission by the mosquito vector, raising the possibility that this reflects evolutionary adaptation of the *P. falciparum* population to the different *Anopheles* species that transmit malaria in different geographical regions.

**Figure 4.**
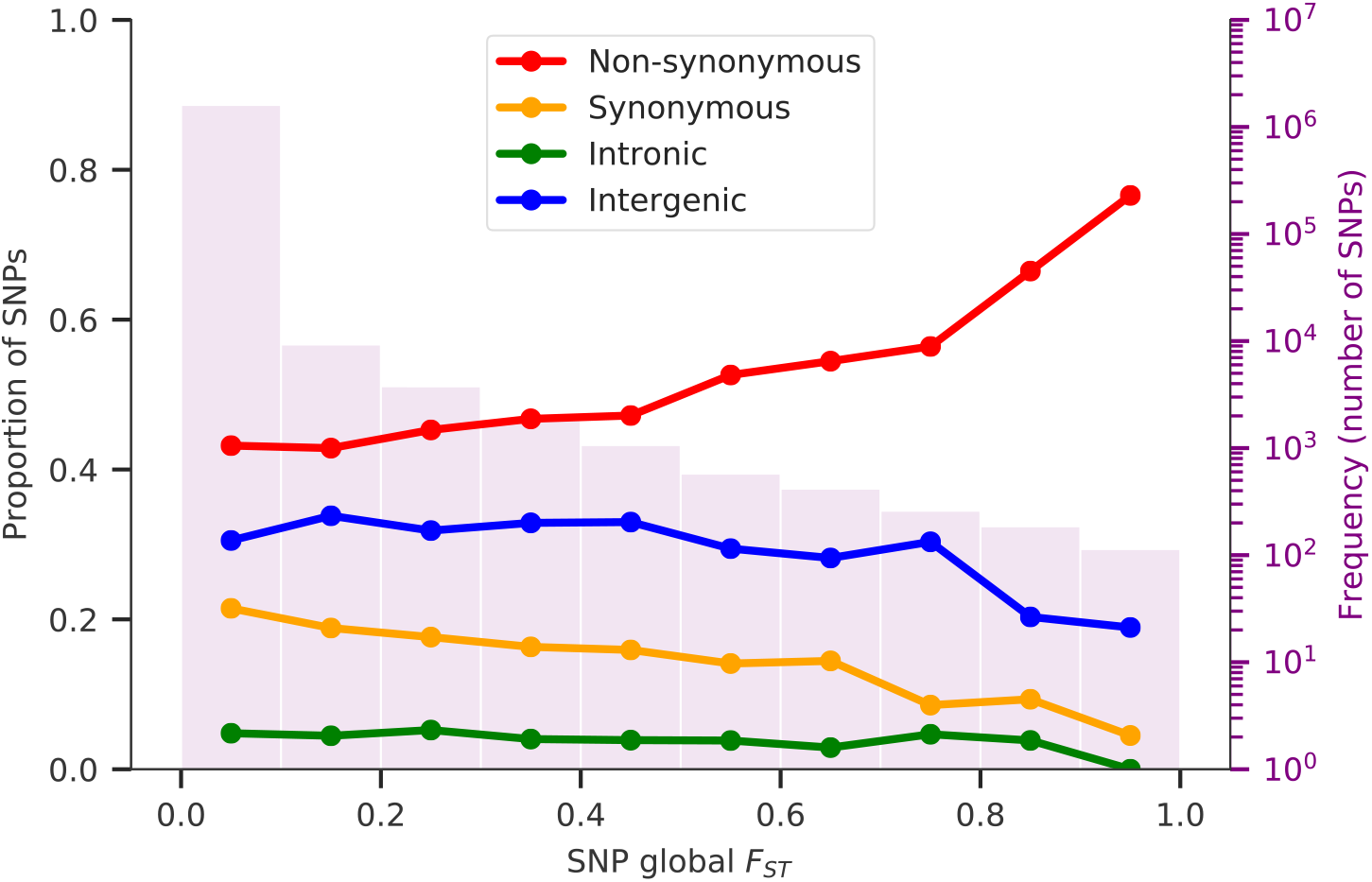
SNPs geographic differentiation. Coloured lines show the proportions of SNPs in ten F_ST_ bins, stratified by genomic regions: non-synonymous (red), synonymous (yellow), intronic (green) and intergenic (blue). F_ST_ is calculated between all eight regional parasite populations and the number of SNPs in each bin is indicated in the background histogram. The y-axis on the right-hand side refers to the histogram and is on a log scale.

It is more difficult to characterise variants that are strongly differentiated at the local level, due to smaller sample sizes and various sources of sampling bias, but a crude estimate can be obtained by analysis of each of the six geographical regions with samples from multiple countries. *F*_*ST*_ was estimated for each SNP across different sampling locations within each geographical region, and the results for different regions were combined by a heuristic approach to obtain a *local differentiation score* for each gene (see Methods). A range of genes associated with drug resistance *(crt*, *dhfr*, *dhps*, *kelch13*, *mdr1*, *mdr2* and *fd*) were in the top centile of local differentiation scores (Supplementary Figure 1, Supplementary Table 8, Supplementary Note).

### Geographic patterns of drug resistance

#### Classification of samples based on markers of drug resistance

Antimalarial drug resistance represents a major focus of research for many partner studies within the *Pf* Community Project, and this dataset therefore contains a significant body of data that have appeared in previous reports on drug resistance. Readers are referred to these publications for more detailed analyses of local patterns of resistance^9–14,16–22^ and of resistance to specific drugs including chloroquine^16,21^, sulfadoxine-pyrimethamine^16,19,21^ and artemisinin combination therapy^9–11,13–15,17,18,21,22^.

Here we have classified all samples into different types of drug resistance based on published genetic markers and current knowledge of the molecular mechanisms (see www.malariagen.net/resource/26 for details of the heuristic used). Table 2 summarises the frequency of different types of drug resistance in samples from different geographical regions. Overall, we observed higher prevalence of samples classified as resistant in Southeast Asia than anywhere else, with multiple samples resistant to all drugs considered. Note that samples were collected over a relatively long time period (2002-15) during which there were major changes in global patterns of drug resistance, and that the sampling locations represented in a given year depended on which partner studies were operative at the time. To alleviate this problem we have also divided the data into samples collected before and after 2011 (Supplementary table 10), but temporal trends in aggregated data should be interpreted with due caution.

**Table 2.**
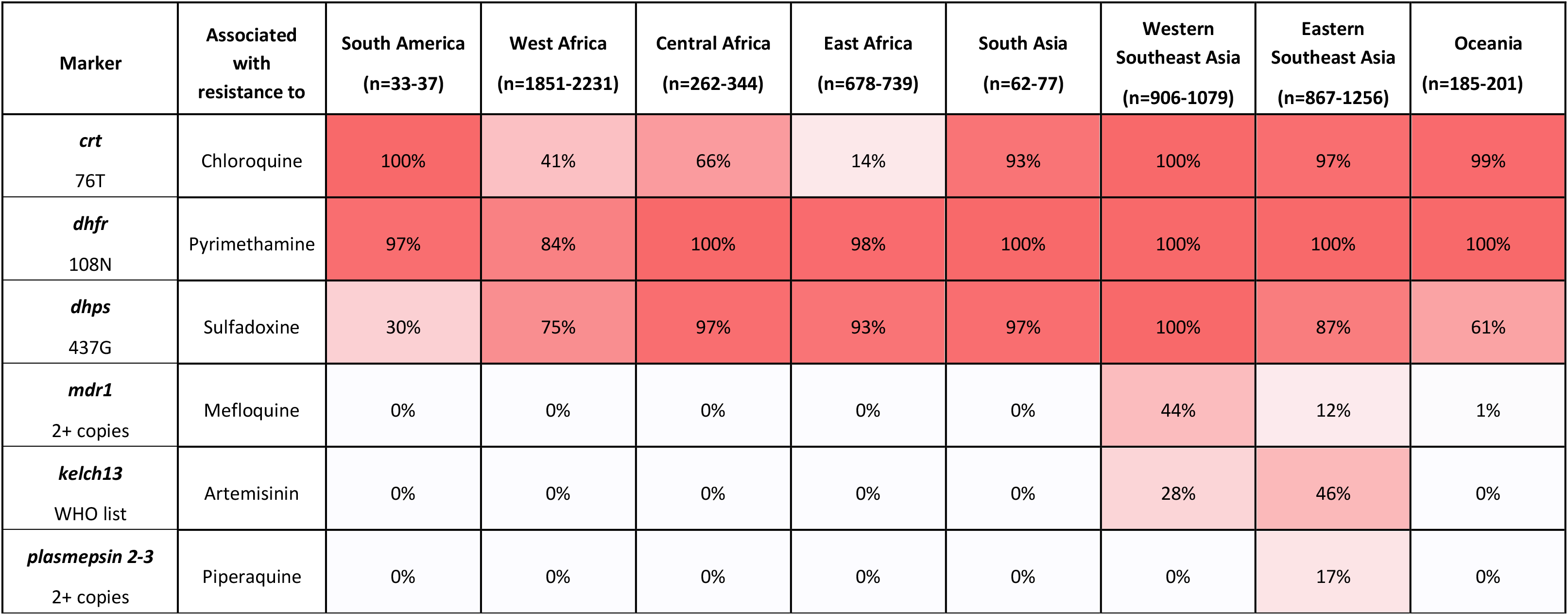
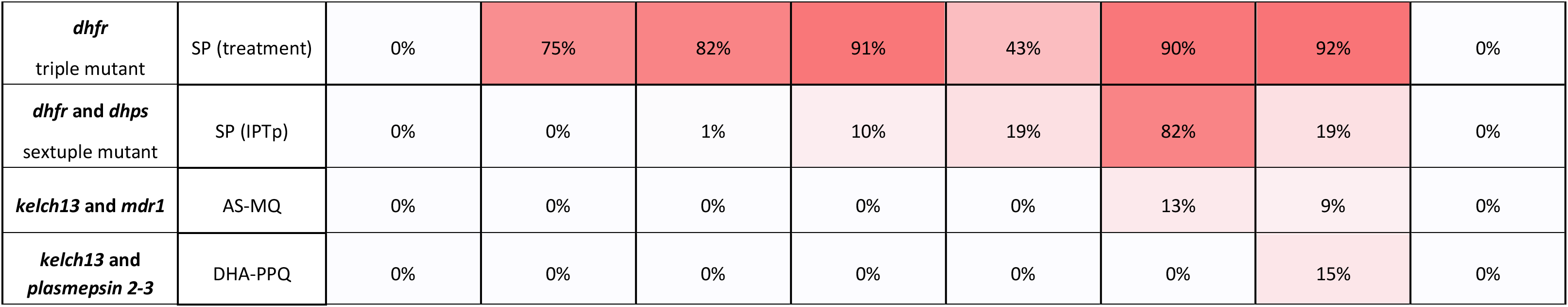
Cumulative frequency of different types of drug resistance in samples from different geographical regions. All samples were classified into different types of drug resistance based on published genetic markers, and represent best attempt based on the available data. Each type of resistance was considered to be either present, absent or unknown for a given sample. For each resistance type, the table reports: the genetic markers considered; the drug they are associated with; the proportion of samples in each region classified as resistant out of the samples where the type was not unknown. The number of samples classified as either resistant or not resistant varies for each type of resistance considered (e.g. due to different levels of genomic accessibility); numbers in brackets reports the minimum and maximum number analysed while the exact numbers considered are reported in Supplementary table 9. SP: sulfadoxine-pyrimethamine; treatment: SP used for the clinical treatment of uncomplicated malaria; IPTp: SP used for intermittent preventive treatment in pregnancy; AS-MQ: artesunate + mefloquine combination therapy; DHA-PPQ: dihydroartemisinin + piperaquine combination therapy. Details of the rules used to infer resistance status from genetic markers can be found on the resource page at www.malariagen.net/resource/26.

| Marker | Associated with resistance to | South America<br>(n=33-37) | West Africa<br>(n=1851-2231) | Central Africa<br>(n=262-344) | East Africa<br>(n=678-739) | South Asia<br>(n=62-77) | Western<br>Southeast Asia<br>(n=906-1079) | Eastern<br>Southeast Asia<br>(n=867-1256) | Oceania<br>(n=185-201) |
| --- | --- | --- | --- | --- | --- | --- | --- | --- | --- |
| *crt*<br>76T | Chloroquine | 100% | 41% | 66% | 14% | 93% | 100% | 97% | 99% |
| *dhfr*<br>108N | Pyrimethamine | 97% | 84% | 100% | 98% | 100% | 100% | 100% | 100% |
| *dhps*<br>437G | Sulfadoxine | 30% | 75% | 97% | 93% | 97% | 100% | 87% | 61% |
| *mdr1*<br>2+ copies | Mefloquine | 0% | 0% | 0% | 0% | 0% | 44% | 12% | 1% |
| *kelch13*<br>WHO list | Artemisinin | 0% | 0% | 0% | 0% | 0% | 28% | 46% | 0% |
| *plasmepsin 2-3*<br>2+ copies | Piperaquine | 0% | 0% | 0% | 0% | 0% | 0% | 17% | 0% |
| ***dhfr***<br>triple mutant | SP (treatment) | 0% | 75% | 82% | 91% | 43% | 90% | 92% | 0% |
| ***dhfr* and *dhps***<br>sextuple mutant | SP (IPTp) | 0% | 0% | 1% | 10% | 19% | 82% | 19% | 0% |
| ***kelch13* and *mdr1*** | AS-MQ | 0% | 0% | 0% | 0% | 0% | 13% | 9% | 0% |
| ***kelch13* and**<br>***plasmepsin 2-3*** | DHA-PPQ | 0% | 0% | 0% | 0% | 0% | 0% | 15% | 0% |

Below we summarise the overall profile of drug resistance types in the regional sub-populations: this is intended simply to provide context for users of this dataset, and should not be regarded as a statement of the current epidemiological situation. The Supplementary Notes contain a more detailed description of the geographical distribution of haplotypes, CNV breakpoints, interactions between genes, and variants associated with less commonly used antimalarial drugs. In the accompanying data release we also identify samples with *mdr1*, *plasmepsin2/3* and *gch1* gene amplifications that can affect drug resistance.

#### Chloroquine resistance

Samples were classified as chloroquine resistant if they carried the *crt* 76T allele. As shown in Table 2, this was found in almost all samples from Southeast Asia, South America and Oceania. It was also found across Africa but at lower frequencies, particularly in East Africa where chloroquine resistance is known to have declined since chloroquine was discontinued^63–65^. Supplementary Table 11 shows the geographical distribution of different *crt* haplotypes (based on amino acid positions 72-76) which is consistent with the theory that chloroquine resistance spread from SE Asia to Africa with multiple independent origins in South America and Oceania^66,67^. The *crt* locus is also relevant to other types of drug resistance, e.g. *crt* variants that are relatively specific to SE Asia form the genetic background of artemisinin resistance, and newly emerging *crt* alleles have been associated with the spread of ACT failure due to piperaquine resistance^13,14,22,68^.

#### Sulfadoxine-pyrimethamine resistance

Clinical resistance to sulfadoxine-pyrimethamine is determined by multiple mutations and their interactions, so following current practice^69^ we classified SP resistant samples into four overlapping types: (i) carrying the dhfr 108N allele, associated with pyrimethamine resistance; (ii) the *dhps* 437G allele, associated with sulfadoxine resistance; (iii) carrying the *dhfr* triple mutant, which is strongly associated with SP failure; (iv) carrying the *dhfr/dhps* sextuple mutant, which confers a higher level of SP resistance. As shown in Table 2, *dhfr* 108N was found in almost all samples in all regions apart from West Africa, while *dhps* 437G was at very high frequency throughout most of Africa and Asia, and at lower frequencies in South America and Oceania (see also Supplementary Table 12). Triple mutant *dhfr* parasites were common throughout Africa and Asia, whereas sextuple mutant *dhfr/dhps* parasites were at much lower frequency except in Western SE Asia. In the accompanying data release we also identify samples with *gch1* gene amplifications (Supplementary Table 4) that can modulate SP resistance^70^, although their effect on the clinical outcome and interaction with mutations in *dhfr* and *dhps* is not fully established.

#### Resistance to artemisinin combination therapy

We classified samples as artemisinin resistant based on the World Health Organisation classification of non-synonymous mutations in the propeller region of the *kelch13* gene that have been associated with delayed parasite clearance^71^. By this definition, artemisinin resistance was confined to Southeast Asia but, as previously reported, this dataset contains a substantial number of non-synonymous *kelch13* propeller SNPs occurring at <5% frequency in Africa and elsewhere^9^. The most common ACT formulations in Southeast Asia are artesunate-mefloquine (AS-MQ) and dihydroartemisinin-piperaquine (DHA-PPQ). We classified samples as mefloquine resistant if they had *mdr1* amplification^72^ or as piperaquine resistant if they had *plasmepsin 2/3* amplification^25^. Mefloquine resistance was observed throughout SE Asia and was most common in the western part. Piperaquine resistance was confined to eastern SE Asia with a notable concentration in western Cambodia. Elsewhere^11,13^ we describe the KEL1/PLA1 lineage of artemisinin- and piperaquine-resistant parasites that expanded in western Cambodia during 2008-13, and then spread to other countries during 2013-18, causing high rates of DHA-PPQ treatment failure across eastern SE Asia: since the current dataset extends only to 2015 it captures only the first phase of the KEL1/PLA1 lineage expansion.

### HRP2/3 deletions that affect rapid diagnostic tests

Rapid diagnostic tests (RDTs) provide a simple and inexpensive way to test for parasites in the blood of patients who are suspected to have malaria, and have become a vital tool for malaria control^73,74^. The most widely used RDTs are designed to detect *P. falciparum* histidine-rich protein 2 and cross-react with histidine-rich protein 3, encoded by the *hrp2* and *hrp3* genes respectively. Parasites with gene deletions of *hrp2* and/or *hrp3* have emerged as an important cause of RDT failure in a number of locations^75–79^. It is difficult to devise a simple genetic assay to monitor for risk of RDT failure because *hrp2* and *hrp3* deletions comprise a diverse mixture of large structural variations with multiple independent origins, and both genes are located in subtelomeric regions of the genome with very high levels of natural variation^29,80–83^. In the absence of a well-validated algorithmic method, we visually inspected sequence read coverage and identified samples with clear evidence of large structural variants that disrupted or deleted the *hrp2* and *hrp3* genes. We took a conservative approach: samples that appeared to have a mixture of deleted and non-deleted genotypes were classified as non-deleted.

Deletions were found at relatively high frequency in Peru (8 of 21 samples had *hrp2* deletions, 14 had *hrp3* deletions and 6 had both) but were not seen in samples from Colombia and were relatively rare outside South America. Oceania was the only other region where we observed *hrp2* deletions, but at very low frequency (4%, n=3/80), and also had *hrp3* deletions (25%) though no combined deletions were seen. Deletions of *hrp3* only were more geographically widespread than *hrp2* deletions, being common in Ethiopia (43%, n=9/21) and in Senegal (7%, n=6/84), and at relatively low frequency (<5%) in Kenya, Cambodia, Laos, and Vietnam (Supplementary Table 13). Note that these findings might under-estimate the true prevalence of *hrp2*/*hrp3* deletions, due to sampling bias (our samples were primarily collected from RDT-positive cases) and also because we focused on large structural variants and did not consider polymorphisms that might also cause RDT failure but would require more sophisticated analytical approaches. There is a need for more reliable diagnostics of *hrp2* and *hrp3* deletions, and we hope that these open data will accelerate this important area of applied methodological research.

## Discussion

This open dataset comprises sequence reads and genotype calls on over 7,000 *P. falciparum* samples from MalariaGEN partner studies in 28 countries. After excluding variants and samples that failed to meet stringent quality control criteria, the dataset contains high-quality genotype calls for 3 million polymorphisms including SNPs, indels, CNVs and large structural variations, in almost 6,000 samples. The data can be analysed in their entirety or can be filtered to select for specific genes, or geographical locations, or samples with particular genotypes. This is twice the sample size of our previous consortial publication^9^ and is the largest available data resource for analysis of *P. falciparum* population structure, gene flow and evolutionary adaptation. Each sample has been annotated to show its profile of resistance to six major antimalarial drugs and whether it carries structural variations that can cause RDT failure. The classification scheme is heuristic and based on a subset of known genetic markers, so it should not be treated as a failsafe predictor of the phenotype of a particular sample. Our purpose in providing these annotations is to make it easy for users without specialist training in genetics to explore the global dataset and to analyse any subset of samples for key features that are relevant to malaria control.

An important function of this curated dataset is to provide information on the provenance and key features of samples associated with each partner study, thus allowing the findings reported in different publications to be linked and compared. Data produced by the *Pf* Community Project have been analysed in more than 50 publications (refs 5-55) and a few examples will serve to illustrate the diverse ways in which the data are being used. An analysis of samples collected across Africa by Amambua-Ngwa, Djimde and colleagues found evidence that parasite population structure overlaps with historical patterns of human migration and that the *P. falciparum* population in Ethiopia is significantly diverged from other parts of the continent.^12^ A series of studies by Amato, Miotto and colleagues have documented the evolution of a multidrug-resistant lineage of *P. falciparum* that originated in Western Cambodia over ten years ago and is now expanding rapidly across Southeast Asia, acquiring additional resistance mutations as it spreads.^11,13,14^ McVean and colleagues have developed a computational method for deconvolution of the haplotypic structure of mixed infections, allowing analysis of the pedigree structure of parasites that are cotransmitted by the same mosquito.^49^ Bahlo and colleagues have developed a different haplotype-based method to describe the relatedness structure of the parasite population and to identify new genomic loci with evidence of recent positive selection.^50^

A recent report from the World Health Organisation highlights the need for improved surveillance systems in sustaining malaria control and achieving the long-term goal of malaria eradication.^84^ To be of practical value for national malaria control programmes, genetic data must address well-defined use cases and be readily accessible.^85^ Amplicon sequencing technologies provide a powerful new tool for targeted genotyping that could feasibly be implemented locally in malaria-endemic countries ^86,87^, but there remains a need for the international malaria control community to generate and share whole genome sequencing data, e.g. to monitor for newly emerging forms of drug resistance and to understand regional patterns of parasite migration. The next generation of long-read sequencing technologies will improve the precision of population genomic inference, e.g. by enabling analysis of hypervariable regions of the genome, and of pedigree structures within mixed infections. The accuracy with which the resistance phenotype of a sample can be predicted from genome sequencing data will also improve as we gain better functional understanding of the polygenic determinants of drug resistance.

Thus the next few years are likely to see major advances in both the scale and information content of parasite genomic data. The practical value for malaria control will be greatly enhanced by the progressive acquisition of longitudinal time-series data, particularly if this is linked to other sources of epidemiological data and translated into reliable, actionable information with sufficient rapidity to allow control programmes to monitor the impact of their interventions on the parasite population in near real time. The *Pf* Community Project provides proof of concept that systems can be developed for groups in different countries to share data, to analyse it using standardised methods, and to make it readily accessible to other researchers and the malaria control community.

## Methods

All samples in this study were derived from blood samples obtained from patients with *P. falciparum* malaria, collected with informed consent from the patient or a parent or guardian. At each location, sample collection was approved by the appropriate local and institutional ethics committees. The following local and institutional committees gave ethical approval for the partner studies: Human Research Ethics Committee of the Northern Territory Department of Health & Families and Menzies School of Health Research, Darwin, Australia; National Research Ethics Committee of Bangladesh Medical Research Council, Bangladesh; Comite d’Ethique de la Recherche - Institut des Sciences Biomedicales Appliquees, Benin; Ministere de la Sante – Republique du Benin, Benin; Comité d’Éthique, Ministère de la Santé, Bobo-Dioulasso, Burkina Faso; Institutional Review Board Centre Muraz, Burkina Faso; Ministry of Health National Ethics Committee for Health Research, Cambodia; Institutional Review Board University of Buea, Cameroon; Comite Institucional de Etica de investigaciones en humanos de CIDEIM, Colombia; Comité National d’Ethique de la Recherche, Cote d’Ivoire; Comite d’Ethique Universite de Kinshasa, Democratic Republic of Congo; Armauer Hansen Research Institute Institutional Review Board, Ethiopia; Addis Ababa University, Aklilu Lemma Institute of Pathobiology Institutional Review Board, Ethiopia; Kintampo Health Research Centre Institutional Ethics Committee, Ghana; Ghana Health Service Ethical Review Committee, Ghana; University of Ghana Noguchi Medical Research Institute, Ghana; Navrongo Health Research Centre Institutional Review Board, Ghana; Comite d’Ethique National Pour la Recherché en Santé, Republique de Guinee; Indian Council of Medical Research, India; Eijkman Institute Research Ethics Commission, Eijkman Institute for Molecular Biology, Jakarta, Indonesia; KEMRI Scientific and Ethics Review Unit, Kenya; Ministry of Health National Ethics Committee For Health Research, Laos; Ethical Review Committee of University of Ilorin Teaching Hospital, Nigeria; Comité National d’Ethique auprès du Ministère de la Santé Publique, Madagascar; College of Medicine Regional Ethics Committee University of Malawi, Malawi; Faculté de Médecine, de Pharmacie et d’Odonto-Stomatologie, University of Bamako, Bamako, Mali; Ethics Committee of the Ministry of Health, Mali; Ethics committee of the Ministry of Health, Mauritania; Department of Medical Research (Lower Myanmar); Ministry of Health, Government of The Republic of the Union of Myanmar;: Institutional Review Board, Papua New Guinea Institute of Medical Research, Goroka, Papua New Guinea; PNG Medical Research Advisory Council (MRAC), Papua New Guinea; Institutional Review Board, Universidad Nacional de la Amazonia Peruana, Iquitos, Peru; Ethics Committee of the Ministry of Health, Senegal; National Institute for Medical Research and Ministry of Health and Social Welfare, Tanzania; Medical Research Coordinating Committee of the National Institute for Medical Research, Tanzania; Ethics Committee, Faculty of Tropical Medicine, Mahidol University, Bangkok, Thailand; Ethics Committee at Institute for the Development of Human Research Protections, Thailand; Gambia Government/MRC Joint Ethics Committee, Banjul, The Gambia; London School of Hygiene and Tropical Medicine Ethics Committee, London, UK; Oxford Tropical Research Ethics Committee, Oxford, UK; Walter Reed Army Institute of Research, USA; National Institute of Allergy and Infectious Diseases, Bethesda, MD, USA; Ethical Committee, Hospital for Tropical Diseases, Ho Chi Minh City, Vietnam; Ministry of Health Institute of Malariology-Parasitology-Entomology, Vietnam.

Standard laboratory protocols were used to determine DNA quantity and proportion of human DNA in each sample as previously described^7,56^.

Here we summarise the bioinformatics methods used to produce and analyse the data; all the details are available at www.malariagen.net/resource/26.

Reads mapping to the human reference genome were discarded before all analyses, and the remaining reads were mapped to the *P. falciparum* 3D7 v3 reference genome using bwa mem^88^. “Improved” BAMs were created using the Picard tools `CleanSam, FixMateInformation` and `MarkDuplicates` and GATK base quality score recalibration. All lanes for each sample were merged to create sample-level BAM files.

We discovered potential SNPs and indels by running GATK’s `HaplotypeCaller`^89^ independently across each of the 7,182 sample-level BAM files and genotyped these for each of the 16 reference sequences (14 chromosomes, 1 apicoplast and 1 mitochondria) using GATK’s `CombineGVCFs` and `GenotypeGCVFs`.

SNPs and indels were filtered using GATK’s Variant Quality Score Recalibration (VQSR). Variants with a VQSLOD score ≤ 0 were filtered out. Functional annotations were applied using snpEff^90^ version 4.1. Genome regions were annotated using `vcftools` and masked if they were outside the core genome. Unless otherwise specified, we used biallelic SNPs that pass all quality filters for all the analysis.

We removed 69 samples from lab studies to create the release VCF files which contain 7,113 samples. VCF files were converted to ZARR format and subsequent analyses were mainly performed using `scikit-allel` (https://github.com/cggh/scikit-allel) and the ZARR files.

We identified species using nucleotide sequence from reads mapping to six different loci in the mitochondrial genome, using custom java code (https://github.com/malariagen/GeneticReportCard). The loci were located within the *cox3* gene (PF3D7_MIT01400), as described in a previously published species detection method.^91^ Alleles at various mitochondrial positions within the six loci were genotyped and used for classification as shown in Supplementary Table 14.

We created a final analysis set of 5,970 samples after removing replicate, low coverage, suspected contaminations or mislabelling and mixed-species samples.

We calculate genetic distance between samples using biallelic SNPs that pass filters using a method previously described^9^. In addition to calculating genetic distance between all pairs of samples from the current data set, we also calculated the genetic distance between each sample and the lab strains 3D7, 7G8, GB4, HB3 and Dd2 from the Pf3k project (www.malariagen.net/projects/pf3k).

The matrix of genetic distances was used to generate neighbour-joining trees and principal coordinates. Based on these observations we grouped the samples into eight geographic regions: South America, West Africa, Central Africa, East Africa, South Asia, the western part of Southeast Asia, the eastern part of Southeast Asia and Oceania, with samples assigned to region based on the geographic location of the sampling site. Five samples from returning travellers were assigned to region based on the reported country of travel.

*F*_*WS*_ was calculated using custom python scripts using the method previously described^7^. Nucleotide diversity (π) was calculated in non-overlapping 25kbp genomic windows, only considering coding biallelic SNPs to reduce the ascertainment bias caused by poor accessibility of non-coding regions. LD decay (*r*^2^) was calculated using the method of Rogers and Huff and biallelic SNPs with low missingness and regional allele frequency >10%. Mean *F*_*ST*_ between populations was calculated using Hudson’s method.

Allele frequencies stratified by geographic regions and sampling sites were calculated using the genotype calls produced by GATK. *F*_*ST*_ was calculated between all 8 regions, and also between all sites with at least 25 QC pass samples. *F*_*ST*_ between different locations for individual SNPs was calculated using Weir and Cockerham’s method.

We used two complementary methods to determine tandem duplication genotypes around *mdr1*, *plasmepsin2/3* and *gch1*, namely a coverage-based method and a method based on position and orientation of reads near discovered duplication breakpoints. In brief, the outline algorithm is: (1) Determine copy number at each locus using a coverage based hidden Markov model (HMM); (2) Determine breakpoints of identified duplications by manual inspection of reads and face-away read pairs around all sets of breakpoints; (3) for each locus in each sample, initially set copy number to that determined by the HMM if <= 10 CNVs discovered in total, else consider undetermined; (4) if face-away pairs provide self-sufficient evidence for the presence or absence of the amplification, override the HMM call; (5) for each locus in each sample, set the breakpoint to be that with the highest proportion of face-away reads.

We genotyped deletions in *hrp2* and *hrp3* by manual inspection of sequence read coverage plots.

The procedure used to map genetic markers to inferred resistance status classification is described in the details for each drug in the accompanying data release (https://www.malariagen.net/resource/26).

In brief, we called amino acids at selected loci by first determining the reference amino acids and then, for each sample, applying all variations using the GT field of the VCF file. The amino acid and copy number calls generated were used to classify all samples into different types of drug resistance. Our methods of classification were heuristic and based on the available data and current knowledge of the molecular mechanisms. Each type of resistance was considered to be either present, absent or unknown for a given sample.

We defined the global differentiation score for a gene as 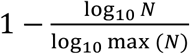, where *N* is the rank of the non-synonymous SNP with the highest global *F*_*ST*_ value within that gene. To define the local differentiation score, we first calculated for each region containing multiple sites (WAF, EAF, SAS, WSEA, ESEA and OCE) *F*_*ST*_ for each SNP between sites within that region. For each gene, we then calculated the rank of the highest *F*_*ST*_ non-synonymous SNP within that gene for each of the six regions. We defined the local differentiation score for each gene using the second highest of these six ranks (N), to ensure that the gene was highly ranked in at least two populations, i.e. to minimise the chance of artefactually ranked a gene highly due to a single variant in a single population. The final local differentiation score was normalised to ensure that the range of possible scores was between 0 and 1, local differentiation score was defined as 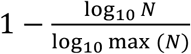.

## Authors

MalariaGEN *Plasmodium falciparum* Community Project

### Data analysis group

Pearson, RD^1, 2, *^, Amato, R^1, 2, *^, Hamilton, WL^1, 3^, Almagro-Garcia, J^1, 2^, Chookajorn, T^4^, Kochakarn, T^1, 4^, Miotto, O^1, 2, 5^, Kwiatkowski, DP^1, 2, 6^

*Joint analysis lead

### Local study design, implementation and sample collection

Ahouidi, A^7^, Amambua-Ngwa, A^8^, Amaratunga, C^9^, Amenga-Etego, L^10, 11^, Andagalu, B^12^, Anderson, TJ^13^, Andrianaranjaka, V^14^, Apinjoh, T^15^, Ashley, E^5^, Auburn, S^16, 17^, Awandare, G^11, 18^, Ba, H^19^, Baraka, V^20, 21^, Barry, AE^22, 23, 24^, Bejon, P^25^, Bertin, GI^26^, Boni, MF^17, 27^, Borrmann, S^28^, Bousema, T^29, 30^, Branch, O^31^, Bull, PC^25, 32^, Chotivanich, K^4^, Claessens, A^8^, Conway, D^29^, Craig, A^33, 34^, d’Alessandro, U^8^, Dama, S^35^, Day, N^5^, Diakite, M^35^, Djimdé, A^35^, Dolecek, C^17^, Dondorp, A^5^, Drakeley, C^29^, Duffy, P^9^, Echeverry, DF^36, 37^, Egwang, TG^38^, Erko, B^39^, Fairhurst, RM^40^, Faiz, A^41^, Fanello, CA^5^, Fukuda, MM^42^, Gamboa, D^43^, Ghansah, A^44^, Golassa, L^39^, Harrison, GLA^24^, Hien, TT^27, 45^, Hill, CA^46^, Hodgson, A^47^, Imwong, M^4^, Ishengoma, DS^20, 48^, Jackson, SA^49^, Kamaliddin, C^26^, Kamau, E^50^, Konaté, A^51^, Kyaw, MP^52, 53^, Lim, P^54, 9^, Lon, C^42^, Loua, KM^55^, Maïga-Ascofaré, O^35, 56, 57^, Marfurt, J^16^, Marsh, K^17, 58^, Mayxay, M^59, 60^, Mobegi, V^61^, Mokuolu, OA^62^, Montgomery, J^63^, Mueller, I^24, 64^, Newton, PN^65^, Nguyen, T^27^, Noedl, H^66^, Nosten, F^17, 67^, Noviyanti, R^68^, Nzila, A^69^, Ochola-Oyier, LI^25^, Ocholla, H^70, 71, 72^, Oduro, A^10^, Onyamboko, MA^73^, Ouedraogo, J^74^, Peshu, N^25^, Phyo, AP^5, 67^, Plowe, CV^75^, Price, RN^16, 45, 5^, Pukrittayakamee, S^4^, Randrianarivelojosia, M^14, 76^, Rayner, JC^1^, Ringwald, P^77^, Ruiz, L^78^, Saunders, D^42^, Shayo, A^79^, Siba, P^80^, Su, X^9^, Sutherland, C^29^, Takala-Harrison, S^81^, Tavul, L^80^, Thathy, V^25^, Tshefu, A^82^, Verra, F^83^, Vinetz, J^43, 84^, Wellems, TE^9^, Wendler, J^6^, White, NJ^5^, Yavo, W^51, 85^, Ye, H^86^

### Sequencing, data production and informatics

Pearson, RD^1, 2^, Stalker, J^1^, Ali, M^1^, Amato, R^1, 2^, Ariani, C^1^, Busby, G^2^, Drury, E^1^, Hart, L^2^, Hubbart, C^6^, Jacob, C^1^, Jeffery, B^2^, Jeffreys, AE^6^, Jyothi, D^1^, Kekre, M^1^, Kluczynski, K^2^, Malangone, C^1^, Manske, M^1^, Miles, A^1, 2^, Nguyen, T^1^, Rowlands, K^6^, Wright, I^2^, Goncalves, S^1^, Rockett, KA^1, 6^

### Partner study support and coordination

Simpson, VJ^2^, Miotto, O^1, 2, 5^, Amato, R^1, 2^, Goncalves, S^1^, Henrichs, C^2^, Johnson, KJ^2^, Pearson, RD^1, 2^, Rockett, KA^1, 6^, Kwiatkowski, DP^1, 2, 6^

### Affiliations

^1^Wellcome Sanger Institute, Hinxton, UK

^2^MRC Centre for Genomics and Global Health, Big Data Institute, University of Oxford, Oxford, UK

^3^Cambridge University Hospitals NHS Foundation Trust, Cambridge, UK

^4^Mahidol University, Bangkok, Thailand

^5^Mahidol-Oxford Tropical Medicine Research Unit (MORU), Thailand

^6^Wellcome Centre for Human Genetics, University of Oxford, Oxford, UK

^7^Hopital Le Dantec, Universite Cheikh Anta Diop, Dakar, Senegal

^8^Medical Research Council Unit, The Gambia

^9^National Institute of Allergy and Infectious Diseases (NIAID), NIH, USA

^10^Navrongo Health Research Centre, Ghana

^11^West African Centre for Cell Biology of Infectious Pathogens (WACCBIP)

^12^United States Army Medical Research Directorate-Africa, Kenya Medical Research Institute/Walter Reed Project, Kisumu, Kenya

^13^Texas Biomedical Research Institute, San Antonio, USA

^14^Institut Pasteur de Madagascar

^15^University of Buea, Cameroon

^16^Menzies School of Health Research, Darwin, Australia

^17^Nuffield Department of Medicine, University of Oxford, UK

^18^University of Ghana, Legon, Ghana

^19^Institut National de Recherche en Santé Publique, Nouakchott, Mauritania

^20^National Institute for Medical Research (NIMR), United Republic of Tanzania

^21^Department of Epidemiology, International Health Unit, Universiteit Antwerpen, Belgium

^22^Deakin University, Australia

^23^Burnet Institute, Australia

^24^Walter and Eliza Hall Institute, Australia

^25^KEMRI Wellcome Trust Research Programme, Kenya

^26^Research institute for development, France

^27^Oxford University Clinical Research Unit (OUCRU), Vietnam

^28^Institute for Tropical Medicine, University of Tübingen, Germany

^29^London School of Hygiene and Tropical Medicine, UK

^30^Radboud University Medical Center, The Netherlands

^31^NYU School of Medicine Langone Medical Center, USA

^32^Department of Pathology, University of Cambridge, UK

^33^Liverpool School of Tropical Medicine, UK

^34^Malawi-Liverpool-Wellcome Trust Clinical Research Programme

^35^Malaria Research and Training Centre, University of Science, Techniques and Technologies of Bamako, Mali

^36^Centro Internacional de Entrenamiento e Investigaciones Médicas - CIDEIM, Cali, Colombia

^37^Universidad Icesi, Cali, Colombia

^38^Biotech Laboratories, Uganda

^39^Aklilu Lemma Institute of Pathobiology, Addis Ababa University, Ethiopia

^40^National Institutes of Health (NIH), USA

^41^Dev Care Foundation, Dhaka, Bangladesh

^42^Department of Immunology and Medicine, US Army Medical Component, Armed Forces Research Institute of Medical Sciences (USAMC-AFRIMS), Bangkok, Thailand

^43^Laboratorio ICEMR-Amazonia, Laboratorios de Investigacion y Desarrollo, Facultad de Ciencias y Filosofia, Universidad Peruana Cayetano Heredia, Lima, Peru

^44^Nogouchi Memorial Institute for Medical Research, Legon-Accra, Ghana

^45^Centre for Tropical Medicine and Global Health, University of Oxford, UK

^46^Department of Entomology, Purdue University, West Lafayette, USA

^47^Ghana Health Service, Ministry of Health, Ghana

^48^East African Consortium for Clinical Research (EACCR), United Republic of Tanzania

^49^Center for Applied Genetic Technologies, University of Georgia, Athens, GA, USA

^50^Walter Reed Army Institute of Research, U.S. Military HIV Research Program, Silver Spring, MD, USA

^51^University Félix Houphouët-Boigny, Côte d’Ivoire

^52^The Myanmar Oxford Clinical Research Unit, University of Oxford, Myanmar

^53^University of Public Health, Yangon, Myanmar

^54^Medical Care Development International, Maryland, USA

^55^Institut National de Santé Publique, Conakry, Republic of Guinea

^56^Bernhard Nocht Institute for Tropical Medicine, Germany

^57^Research in Tropical Medicine, Kwame Nkrumah University of Sciences and Technology, Kumasi, Ghana

^58^African Academy of Sciences, Kenya

^59^Wellcome Trust-Mahosot Hospital Oxford University Medicine Research Collaboration (LOMWRU)

^60^Faculty of Postgraduate Studies, University of Health Sciences (UHS), Vientiane, Laos

^61^School of Medicine, University of Nairobi, Kenya

^62^Department of Paediatrics and Child Health, University of Ilorin, Ilorin, Nigeria

^63^Institute of Vector-Borne Disease, Monash University, Clayton, Victoria, 3800, Australia

^64^Barcelona Centre for International Health Research, Spain

^65^Wellcome Trust-Mahosot Hospital-Oxford Tropical Medicine Research Collaboration, Lao, PDR

^66^Malaria Research Initiative Bandarban (MARIB), Bangladesh

^67^Shoklo Malaria Research Unit

^68^Eijkman Institute for Molecular Biology, Indonesia

^69^King Fahid University of Petroleum and Minerals (KFUMP), Saudi Arabia

^70^Malaria Capacity Development Consortium

^71^KEMRI - Centres for Disease Control and Prevention (CDC) Research Program, Kisumu, Kenya

^72^Centre for Bioinformatics and Biotechnology, University of Nairobi, Kenya

^73^Kinshasa School of Public Health, University of Kinshasa, DRC

^74^Institut de Recherche en Sciences de la Santé, Burkina Faso

^75^Duke Global Health Institute, Duke University

^76^Universités d’Antananarivo et de Mahajanga, Madagascar

^77^World Health Organization (WHO), Switzerland

^78^Universidad Nacional de la Amazonia Peruana, Peru

^79^Nelson Mandela Institute of Science and Technology, Tanzania

^80^Papua New Guinea Institute of Medical Research, PNG

^81^Center for Vaccine Development and Global Health, University of Maryland School of Medicine, Baltimore, MD, USA

^82^University of Kinshasa, DRC

^83^Sapienza University of Rome, Italy

^84^Yale School of Medicine, New Haven, CT

^85^Malaria Research and Control Center of the National Institute of Public Health, Côte d’Ivoire

^86^Department of Medical Research, Myanmar

## Acknowledgements

This study was conducted by the MalariaGEN *Plasmodium falciparum* Community Project, and was made possible by clinical parasite samples contributed by partner studies, whose investigators are represented in the author list and in the associated data release (https://www.malariagen.net/resource/26). In addition, the authors would like to thank the following individuals who contributed to partner studies, making this study possible: Dr Eugene Laman for work in sample collection in the Republic of Guinea; Dr Abderahmane Tandia and Dr Yacine Deh for work in sample collection in Mauritania; Dr Ibrahim Sanogo for work in sample collection in Mali. Genome sequencing was undertaken by the Wellcome Sanger Institute and we thank the staff of the Wellcome Sanger Institute Sample Logistics, Sequencing, and Informatics facilities for their contribution. The sequencing, analysis, informatics and management of the Community Project are supported by Wellcome through Sanger Institute core funding (098051), a Strategic Award (090770/Z/09/Z) and the Wellcome Centre for Human Genetics core funding (203141/Z/16/Z), and by the MRC Centre for Genomics and Global Health which is jointly funded by the Medical Research Council and the Department for International Development (DFID) (G0600718; M006212). The views expressed here are solely those of the authors and do not reflect the views, policies or positions of the U.S. Government or Department of Defense. Material has been reviewed by the Walter Reed Army Institute of Research. There is no objection to its presentation and/or publication. The opinions or assertions contained herein are the private views of the author, and are not to be construed as official, or as reflecting true views of the Department of the Army or the Department of Defense. The investigators have adhered to the policies for protection of human subjects as prescribed in AR 70–25.

## References

1 Malaria Genomic Epidemiology Network. A global network for investigating the genomic epidemiology of malaria. Nature 2008; 456: 732–7.

2 Chokshi DA, Parker M, Kwiatkowski DP. Data sharing and intellectual property in a genomic epidemiology network: policies for large-scale research collaboration. Bull World Heal Organ 2006; 84: 382–7.

3 Parker M, Bull SJ, de Vries J, Agbenyega T, Doumbo OK, Kwiatkowski DP. Ethical data release in genome-wide association studies in developing countries. PLoS Med 2009; 6: e1000143.

4 Ghansah A, Amenga-Etego L, Amambua-Ngwa A, et al. Monitoring parasite diversity for malaria elimination in sub-Saharan Africa. Science 2014; 345: 1297–8.

5 Auburn S, Campino S, Clark TG, et al. An effective method to purify Plasmodium falciparum DNA directly from clinical blood samples for whole genome high-throughput sequencing. PLoS One 2011; 6: e22213.

6 Venkatesan M, Amaratunga C, Campino S, et al. Using CF11 cellulose columns to inexpensively and effectively remove human DNA from Plasmodium falciparum-infected whole blood samples. Malar J 2012; 11: 41.

7 Manske M, Miotto O, Campino S, et al. Analysis of Plasmodium falciparum diversity in natural infections by deep sequencing. Nature 2012; 487: 375–9.

8 Vauterin P, Jeffery B, Miles A, et al. Panoptes: Web-based exploration of large scale genome variation data. Bioinformatics 2017; 33. DOI:10.1093/bioinformatics/btx410.

9 MalariaGEN Plasmodium falciparum Community Project. Genomic epidemiology of artemisinin resistant malaria. Elife 2016; 5. DOI:10.7554/eLife.08714.

10 Miotto O, Almagro-Garcia J, Manske M, et al. Multiple populations of artemisinin-resistant Plasmodium falciparum in Cambodia. Nat Genet 2013; 45: 648–55.

11 Amato R, Pearson RD, Almagro-Garcia J, et al. Origins of the current outbreak of multidrug-resistant malaria in southeast Asia: a retrospective genetic study. Lancet Infect Dis 2018; 18: 337–45.

12 Amambua-Ngwa A, Amenga-Etego L, Kamau E, et al. Major subpopulations of Plasmodium falciparum in sub-Saharan Africa. Science (80-) 2019; 365: 813–6.

13 Hamilton WL, Amato R, van der Pluijm RW, et al. Evolution and expansion of multidrug-resistant malaria in southeast Asia: a genomic epidemiology study. Lancet Infect Dis 2019; 19: 943–51.

14 van der Pluijm RW, Imwong M, Chau NH, et al. Determinants of dihydroartemisinin-piperaquine treatment failure in Plasmodium falciparum malaria in Cambodia, Thailand, and Vietnam: a prospective clinical, pharmacological, and genetic study. Lancet Infect Dis 2019; 19: 952–61.

15 Ariey F, Witkowski B, Amaratunga C, et al. A molecular marker of artemisinin-resistant Plasmodium falciparum malaria. Nature 2014; 505: 50–5.

16 Nwakanma DC, Duffy CW, Amambua-Ngwa A, et al. Changes in malaria parasite drug resistance in an endemic population over a 25-year period with resulting genomic evidence of selection. J Infect Dis 2014; 209: 1126–35.

17 Ashley EA, Dhorda M, Fairhurst RM, et al. Spread of Artemisinin Resistance in Plasmodium falciparum Malaria. N Engl J Med 2014; 371: 411–23.

18 Kamau E, Campino S, Amenga-Etego L, et al. K13-propeller polymorphisms in Plasmodium falciparum parasites from sub-Saharan Africa. J Infect Dis 2014; 211: 1352–5.

19 Ravenhall M, Benavente ED, Mipando M, et al. Characterizing the impact of sustained sulfadoxine/pyrimethamine use upon the Plasmodium falciparum population in Malawi. Malar J 2016; 15: 575.

20 Gomes AR, Ravenhall M, Benavente ED, et al. Genetic diversity of next generation antimalarial targets: A baseline for drug resistance surveillance programmes. Int J Parasitol Drugs Drug Resist 2017; 7: 174–80.

21 Apinjoh TO, Mugri RN, Miotto O, et al. Molecular markers for artemisinin and partner drug resistance in natural Plasmodium falciparum populations following increased insecticide treated net coverage along the slope of mount Cameroon: Cross-sectional study. Infect Dis Poverty 2017; 6. DOI:10.1186/s40249-017-0350-y.

22 Ross LS, Dhingra SK, Mok S, et al. Emerging Southeast Asian PfCRT mutations confer Plasmodium falciparum resistance to the first-line antimalarial piperaquine. Nat Commun 2018; 9: 3314.

23 Miotto O, Amato R, Ashley EA, et al. Genetic architecture of artemisinin-resistant Plasmodium falciparum. Nat Genet 2015; 47: 226–34.

24 Takala-Harrison S, Jacob CG, Arze C, et al. Independent Emergence of Artemisinin Resistance Mutations Among Plasmodium falciparum in Southeast Asia. J Infect Dis 2014; 211: 670–9.

25 Amato R, Lim P, Miotto O, et al. Genetic markers associated with dihydroartemisinin-piperaquine failure in Plasmodium falciparum malaria in Cambodia: a genotype-phenotype association study. Lancet Infect Dis 2017; 17: 164–73.

26 Borrmann S, Straimer J, Mwai L, et al. Genome-wide screen identifies new candidate genes associated with artemisinin susceptibility in Plasmodium falciparum in Kenya. Sci Rep 2013; 3: 3318.

27 Wendler JP, Okombo J, Amato R, et al. A Genome Wide Association Study of Plasmodium falciparum Susceptibility to 22 Antimalarial Drugs in Kenya. PLoS One 2014; 9: e96486.

28 Zhu L, Tripathi J, Rocamora FM, et al. The origins of malaria artemisinin resistance defined by a genetic and transcriptomic background. Nat Commun 2018; 9: 5158.

29 Sepúlveda N, Phelan J, Diez-Benavente E, et al. Global analysis of Plasmodium falciparum histidine-rich protein-2 (pfhrp2) and pfhrp3 gene deletions using whole-genome sequencing data and meta-analysis. Infect Genet Evol 2018; 62: 211–9.

30 Williams AR, Douglas AD, Miura K, et al. Enhancing blockade of Plasmodium falciparum erythrocyte invasion: assessing combinations of antibodies against PfRH5 and other merozoite antigens. PLoS Pathog 2012; 8: e1002991.

31 Benavente ED, Oresegun DR, de Sessions PF, et al. Global genetic diversity of var2csa in Plasmodium falciparum with implications for malaria in pregnancy and vaccine development. Sci Rep 2018; 8: 15429.

32 Amambua-Ngwa A, Tetteh KK a, Manske M, et al. Population genomic scan for candidate signatures of balancing selection to guide antigen characterization in malaria parasites. PLoS Genet 2012; 8: e1002992.

33 Campino S, Marin-Menendez A, Kemp A, et al. A forward genetic screen reveals a primary role for Plasmodium falciparum Reticulocyte Binding Protein Homologue 2a and 2b in determining alternative erythrocyte invasion pathways. PLOS Pathog 2018; 14: e1007436.

34 Crosnier C, Iqbal Z, Knuepfer E, et al. Binding of Plasmodium falciparum merozoite surface proteins DBLMSP and DBLMSP2 to human immunoglobulin M is conserved amongst broadly diverged sequence variants. J Biol Chem 2016; epub ahead. DOI:10.1074/jbc.M116.722074.

35 Amambua-Ngwa A, Jeffries D, Amato R, et al. Consistent signatures of selection from genomic analysis of pairs of temporal and spatial Plasmodium falciparum populations from the Gambia. Sci Rep 2018; 8. DOI:10.1038/s41598-018-28017-5.

36 Duffy CW, Amambua-Ngwa A, Ahouidi AD, et al. Multi-population genomic analysis of malaria parasites indicates local selection and differentiation at the gdv1 locus regulating sexual development. Sci Rep 2018; 8: 15763.

37 Duffy CW, Ba H, Assefa S, et al. Population genetic structure and adaptation of malaria parasites on the edge of endemic distribution. Mol Ecol 2017; 26: 2880–94.

38 Duffy CW, Assefa SA, Abugri J, et al. Comparison of genomic signatures of selection on Plasmodium falciparum between different regions of a country with high malaria endemicity. BMC Genomics 2015; 16: 527.

39 Mobegi VA, Duffy CW, Amambua-Ngwa A, et al. Genome-wide analysis of selection on the malaria parasite plasmodium falciparum in West African populations of differing infection endemicity. Mol Biol Evol 2014; 31: 1490–9.

40 Shetty AC, Jacob CG, Huang F, et al. Genomic structure and diversity of Plasmodium falciparum in Southeast Asia reveal recent parasite migration patterns. Nat Commun 2019; 10: 2665.

41 Auburn S, Campino S, Miotto O, et al. Characterization of within-host Plasmodium falciparum diversity using next-generation sequence data. PLoS One 2012; 7: e32891.

42 Assefa SA, Preston MD, Campino S, Ocholla H, Sutherland CJ, Clark TG. estMOI: estimating multiplicity of infection using parasite deep sequencing data. Bioinformatics 2014; 30: 1292–4.

43 Murray L, Mobegi VA, Duffy CW, et al. Microsatellite genotyping and genome-wide single nucleotide polymorphism-based indices of Plasmodium falciparum diversity within clinical infections. Malar J 2016; 15: 275.

44 Chang H-H, Worby CJ, Yeka A, et al. THE REAL McCOIL: A method for the concurrent estimation of the complexity of infection and SNP allele frequency for malaria parasites. PLOS Comput Biol 2017; 13: e1005348.

45 O’Brien JD, Iqbal Z, Wendler J, Amenga-Etego L. Inferring Strain Mixture within Clinical Plasmodium falciparum Isolates from Genomic Sequence Data. PLOS Comput Biol 2016; 12: e1004824.

46 Robinson T, Campino SG, Auburn S, et al. Drug-resistant genotypes and multi-clonality in Plasmodium falciparum analysed by direct genome sequencing from peripheral blood of malaria patients. PLoS One 2011; 6: in press.

47 O’Brien JD, Amenga-Etego L, Li R. Approaches to estimating inbreeding coefficients in clinical isolates of Plasmodium falciparum from genomic sequence data. Malar J 2016; 15: 473.

48 Zhu SJ, Almagro-Garcia J, McVean G. Deconvolution of multiple infections in Plasmodium falciparum from high throughput sequencing data. Bioinformatics 2018; 34: 9–15.

49 Zhu SJ, Hendry JA, Almagro-Garcia J, et al. The origins and relatedness structure of mixed infections vary with local prevalence of P. falciparum malaria. Elife 2019; 8. DOI:10.7554/eLife.40845.

50 Henden L, Lee S, Mueller I, Barry A, Bahlo M. Identity-by-descent analyses for measuring population dynamics and selection in recombining pathogens. PLOS Genet 2018; 14: e1007279.

51 Schaffner SF, Taylor AR, Wong W, Wirth DF, Neafsey DE. hmmIBD: software to infer pairwise identity by descent between haploid genotypes. Malar J 2018; 17: 196.

52 Samad H, Coll F, Preston MD, Ocholla H, Fairhurst RM, Clark TG. Imputation-Based Population Genetics Analysis of Plasmodium falciparum Malaria Parasites. PLoS Genet 2015; 11: e1005131.

53 Ravenhall M, Campino S, Clark TG. SV-Pop: population-based structural variant analysis and visualization. BMC Bioinformatics 2019; 20: 136.

54 Jacob CG, Tan JC, Miller BA, et al. A microarray platform and novel SNP calling algorithm to evaluate Plasmodium falciparum field samples of low DNA quantity. BMC Genomics 2014; 15: 719.

55 Preston MD, Assefa S a, Ocholla H, et al. PlasmoView: A Web-based Resource to Visualise Global Plasmodium falciparum Genomic Variation. J Infect Dis 2014; 209: 1808–15.

56 Miles A, Iqbal Z, Vauterin P, et al. Indels, structural variation, and recombination drive genomic diversity in Plasmodium falciparum. Genome Res 2016; 26: 1288–99.

57 Hamilton WL, Claessens A, Otto TD, et al. Extreme mutation bias and high AT content in Plasmodium falciparum. Nucleic Acids Res 2016; 45: 1889–901.

58 Carvalho CMB, Ramocki MB, Pehlivan D, et al. Inverted genomic segments and complex triplication rearrangements are mediated by inverted repeats in the human genome. Nat Genet 2011; 43: 1074–81.

59 Molina-Cruz A, Garver LS, Alabaster A, et al. The human malaria parasite Pfs47 gene mediates evasion of the mosquito immune system. Science (80-) 2013; 340: 984–7.

60 Gardiner DL, Dixon MWA, Spielmann T, et al. Implication of a Plasmodium falciparum gene in the switch between asexual reproduction and gametocytogenesis. Mol. Biochem. Parasitol. 2005; 140: 153–60.

61 Moelans IIMD, Meis JFGM, Kocken C, Konings RNH, Schoenmakers JGG. A novel protein antigen of the malaria parasite Plasmodium falciparum, located on the surface of gametes and sporozoites. Mol Biochem Parasitol 1991; 45: 193–204.

62 Dessens JT, Beetsma AL, Dimopoulos G, et al. CTRP is essential for mosquito infection by malaria ookinetes. EMBO J 1999; 18: 6221–7.

63 Laufer MK, Thesing PC, Eddington ND, et al. Return of Chloroquine Antimalarial Efficacy in Malawi. N Engl J Med 2006; 355: 1959–66.

64 Laufer MK, Takala-Harrison S, Dzinjalamala FK, Stine OC, Taylor TE, Plowe CV. Return of Chloroquine-Susceptible Falciparum Malaria in Malawi Was a Reexpansion of Diverse Susceptible Parasites. J Infect Dis 2010; 202: 801–8.

65 Frosch AEP, Laufer MK, Mathanga DP, et al. Return of Widespread Chloroquine-Sensitive Plasmodium falciparum to Malawi. J Infect Dis 2014; 210: 1110–4.

66 Wootton JC, Feng X, Ferdig MT, et al. Genetic diversity and chloroquine selective sweeps in Plasmodium falciparum. Nature 2002; 418: 320–3.

67 Mita T, Tanabe K, Kita K. Spread and evolution of Plasmodium falciparum drug resistance. Elsevier, 2009.

68 Agrawal S, Moser KA, Morton L, et al. Association of a Novel Mutation in the Plasmodium falciparum Chloroquine Resistance Transporter With Decreased Piperaquine Sensitivity. J Infect Dis 2017; 216: 468–76.

69 Naidoo I, Roper C. Mapping ‘partially resistant’, ‘fully resistant’, and ‘super resistant-malaria. Trends Parasitol 2013; 29: 505–15.

70 Heinberg A, Kirkman L. The molecular basis of antifolate resistance in *Plasmodium falciparum*: looking beyond point mutations. Ann N Y Acad Sci 2015; 1342: 10–8.

71 World Health Organization. Artemisinin and artemisinin-based combination therapy resistance: status report. 2018 https://apps.who.int/iris/handle/10665/274362.

72 Price RN, Uhlemann AC, Brockman A, et al. Mefloquine resistance in Plasmodium falciparum and increased pfmdr1 gene copy number. Lancet. 2004; 364: 438–47.

73 Cheng Q, Gatton ML, Barnwell J, et al. Plasmodium falciparum parasites lacking histidine-rich protein 2 and 3: a review and recommendations for accurate reporting. Malar J 2014; 13: 283.

74 WHO | Malaria rapid diagnostic test performance. Results of WHO product testing of malaria RDTs: round 8 (2016-2018). WHO 2018. https://www.who.int/malaria/publications/atoz/9789241514965/en/ (accessed Aug 22, 2019).

75 Gamboa D, Ho M-F, Bendezu J, et al. A Large Proportion of P. falciparum Isolates in the Amazon Region of Peru Lack pfhrp2 and pfhrp3: Implications for Malaria Rapid Diagnostic Tests. PLoS One 2010; 5: e8091.

76 Rachid Viana GM, Akinyi Okoth S, Silva-Flannery L, et al. Histidine-rich protein 2 (pfhrp2) and pfhrp3 gene deletions in Plasmodium falciparum isolates from select sites in Brazil and Bolivia. PLoS One 2017; 12: e0171150.

77 Parr JB, Verity R, Doctor SM, et al. *Pfhrp2* -deleted *Plasmodium falciparum* parasites in the Democratic Republic of Congo: A national cross-sectional survey. J Infect Dis 2016; 216: jiw538.

78 Menegon M, L’Episcopia M, Nurahmed AM, Talha AA, Nour BYM, Severini C. Identification of Plasmodium falciparum isolates lacking histidine-rich protein 2 and 3 in Eritrea. Infect Genet Evol 2017; 55: 131–4.

79 Bharti PK, Chandel HS, Ahmad A, Krishna S, Udhayakumar V, Singh N. Prevalence of pfhrp2 and/or pfhrp3 Gene Deletion in Plasmodium falciparum Population in Eight Highly Endemic States in India. PLoS One 2016; 11: e0157949.

80 Baker J, Ho M-F, Pelecanos A, et al. Global sequence variation in the histidine-rich proteins 2 and 3 of Plasmodium falciparum: implications for the performance of malaria rapid diagnostic tests. Malar J 2010; 9: 129.

81 Akinyi S, Hayden T, Gamboa D, et al. Multiple genetic origins of histidine-rich protein 2 gene deletion in Plasmodium falciparum parasites from Peru. Sci Rep 2013; 3: 2797.

82 Akinyi Okoth S, Abdallah JF, Ceron N, et al. Variation in Plasmodium falciparum Histidine-Rich Protein 2 (Pfhrp2) and Plasmodium falciparum Histidine-Rich Protein 3 (Pfhrp3) Gene Deletions in Guyana and Suriname. PLoS One 2015; 10: e0126805.

83 Parr JB, Anderson O, Juliano JJ, Meshnick SR. Streamlined, PCR-based testing for pfhrp2- and pfhrp3-negative Plasmodium falciparum. Malar J 2018; 17: 137.

84 WHO Strategic Advisory Group on Malaria Eradication. Malaria eradication: benefits, future scenarios and feasibility. Executive Summary. Geneva: World Health Organisation, 2019.

85 Dalmat R, Naughton B, Kwan-Gett TS, Slyker J, Stuckey EM. Use cases for genetic epidemiology in malaria elimination. Malar J 2019; 18: 163.

86 Early AM, Daniels RF, Farrell TM, et al. Detection of low-density Plasmodium falciparum infections using amplicon deep sequencing. Malar J 2019; 18: 219.

87 Boyce RM, Hathaway N, Fulton T, et al. Reuse of malaria rapid diagnostic tests for amplicon deep sequencing to estimate Plasmodium falciparum transmission intensity in western Uganda. Sci Rep 2018; 8: 10159.

88 Li H, Durbin R. Fast and accurate short read alignment with Burrows-Wheeler transform. Bioinformatics 2009; 25: 1754–60.

89 Depristo MA, Banks E, Poplin R, et al. A framework for variation discovery and genotyping using next-generation DNA sequencing data. Nat Genet 2011; 43: 491–501.

90 Cingolani P, Platts A, Wang LL, et al. A program for annotating and predicting the effects of single nucleotide polymorphisms, SnpEff: SNPs in the genome of Drosophila melanogaster strain w1118; iso-2; iso-3. Fly (Austin); 6: 80–92.

91 Echeverry DF, Deason NA, Davidson J, et al. Human malaria diagnosis using a single-step direct-PCR based on the Plasmodium cytochrome oxidase III gene. Malar J 2016; 15: 128.

