## Supplementary for "An open dataset of *Plasmodium falciparum* genome variation in 7,000 worldwide samples"

MalariaGEN *Plasmodium falciparum* Community Project

#### Supplementary Note

##### Analysis of local differentiation score

The ten genes with highest local differentiation scores are shown in Supplementary Table 8, and differentiation scores for all genes are available in the data release (https://www.malariagen.net/resource/26).

We identified genes in the top centile of differentiation scores that have previously been implicated in drug resistance, but for which a second gene is located nearby on the same chromosome and has a higher local differentiation score. The only example of this we found were the genes PF3D7_1012700 (*nif4*, aka *pph*) and PF3D7_1012900 (*atg18*) which contain SNPs that have been associated with artemisinin resistance[1,2]. *Atg18* has the third highest local differentiation score (0.85) whereas *nif4* is ranked 39$^{th}$ (0.68). Three different SNPs in *nif4*: V1157L, Y1133N and N659S, are the most highly differentiated in WSEA, ESEA and OCE, respectively. For each of these regions, the T38I mutation in *atg18* is more highly differentiated. This lends weight to the hypothesis that *atg18*:T38I is more likely to be the mutation driving the peak seen in GWAS studies[2]

We also identified transporter genes that had high local differentiation scores but which have not to our knowledge previously been directly implicated in drug resistance in *Plasmodium*. Examples include PF3D7_1218400 (local differentiation score 0.76), a phosphate transporter gene with a highly differentiated SNP adjacent to one of the predicted transmembrane domains. The amino acid transporter *aat1* (PF3D7_0629500; local differentiation score 0.71) also contains several SNPs that are highly differentiated in this analysis and is located in a locus that has been candidate before[3,4]. The genomic region has also been previously associated with chloroquine resistance in a GWAS from the China-Myanmar border[2] and variants in an orthologous gene induces chloroquine resistance in yeast cells[5] and in a malaria rodent model[6]. Furthermore, variants have recently been associated with drug resistance in a chemogenomics study[7]. Other transporter genes with high local differentiation scores that have not previously been associated with drug resistance include PF3D7_1440800 (*mfs6*, local differentiation score 0.73) and PF3D7_1129900 (*mfr5*, local differentiation score 0.71).

We also examined which SNPs were driving high local differentiation scores in known drug resistance genes. For example, the most commonly reported mutations in *dhps* are 437G and 540E, but the variants driving the high local differentiation score in *dhps* are 581G (which is the most highly differentiated *dhps* SNP in EAF, WSEA ESEA) and 431V. To date there have been relatively few reports of frequencies of 431V[8–10]. Likewise, the most highly differentiated SNPs in *crt* do not include any in amino acid positions 72-76. These findings highlight the need for constant evaluation of, and monitoring of the changes in allele frequencies of, SNPs in key drug resistance genes.

#### The classic 76T chloroquine resistance mutation in *crt* is found on multiple haplotypes

We analysed the haplotypes at amino acids 72-76 in *crt* (Supplementary table 11). The two most common haplotypes are the wild-type CVMNK which has high frequency in Africa but is rare in Asia, and CV**IET** which is dominant in Asia but also has appreciable frequency across Africa. However, we observe overall seven different *crt* amino acid 72-76 haplotypes. It is worth noting that one haplotype in particular, CV**IDT**, is present at high frequency in ESEA only, and sympatrically with the more common and wide-spread CV**IET**. This high prevalence raises questions about its phenotypic and fitness effects. The amino acid haplotype **S**VMN**T** is dominant in OCE, but also relatively common in SAM. This is two different haplotypes at the nucleotide level with 72S in OCE being due exclusively to a T/A mutation at Pf3D7_07_v3:403,612, whereas the 72S in SAM is due exclusively to a G/C mutation at Pf3D7_07_v3:403,613. Haplotypes CVM**ET** and CVMN**T** are seen exclusively in SAM and **YVIET** is seen exclusively in ESEA.

#### Suplhadoxine-pyrimethamine resistance is widespread and associated with many haplotypes

Many studies on resistance to sulphadoxine-pyrimethamine (S-P) have focused on a small number of specific haplotypes at eight amino acids in the genes *dhfr* (amino acids 51, 59, 108 and 164) and *dhps* (amino acids 437, 540, 581 and 613), though in our dataset we see 62 different haplotypes for these positions (Supplementary Table 12). Most samples have at least four mutations in these codons, with the exception of SAM where the majority of samples have the single 108N mutation (giving the eight amino acid haplotype NC**N**I/AKAA). The majority of samples from WAF and CAF have the quadruple **IRN**I/**G**KAA haplotype, whereas the quintuple haplotype **IRN**I/**GE**AA dominates in EAF. **IRN**I/**GE**AA is also at high frequency throughout Asia, though other quintuple and sextuple mutants are common such as **IRNL**/**GE**AA which is seen in SAS (18%), WSEA (13%) and ESEA (9%). The most common haplotype in both SAS and OCE is the quadruple mutant N**RN**I/**GE**AA, which is relatively rare elsewhere. The most common haplotype in WSEA is the septuple mutant **IRNL**/**GEG**A (58%), whereas the most common in ESEA is the septuple mutant **IRNL**/**GNG**A (22%).

Duplications of *gch1* have also been associated with resistance to S-P. We found eleven different sets of duplication breakpoints around *gch1*, including two examples of DUP-TRP/INV-DUP rearrangements (Supplementary Table 4). DUP-TRP/INV-DUP rearrangements have previously been observed in human data, but to the best of our knowledge this is the first report in Plasmodium species[11]. Most of the common sets of duplication breakpoints were seen across multiple sites and among samples with different *dhfr*/*dhps* haplotypes. An interesting exception to this was the DUP-TRP/INV-DUP rearrangement PfGCH1_DTD_2, which was seen almost exclusively in samples from the island of Papua, all of which carried the N**RN**I/**GE**AA haplotype.

#### *mdr1* duplications have many different breakpoints

Amplifications of the gene *mdr1* are markers of resistance to mefloquine. We identified 28 different sets of breakpoints for *mdr1* duplications (Supplementary Table 5). 27 of these are tandem duplications, but we also see evidence of a DUP-TRP/INV-DUP rearrangement. Many of the more common sets of *mdr1* duplication breakpoints are seen at multiple geographical sites, and also in combination with multiple different *kelch13* mutations, suggesting either gene flow between sites or multiple independent events. Many of the breakpoint pairs share one breakpoint with at least one other pair, suggesting there are breakpoint hotspots around *mdr1*.

#### Artemisinin, piperaquine, and mefloquine resistance

Amplifications of the genes *plasmepsin 2-3* are marker of resistance to piperaquine, a drug commonly used in ACTs[12,13]. We see just three sets of tandem duplication breakpoints (Supplementary Table 5). The 9kb duplication PfPlasmepsin_1 is by far the most common. This is seen at many different sites in ESEA, and is the duplication seen in the strain recently reported to be spreading throughout ESEA[14,15]. The vast majority of samples carrying this duplication also have *kelch13* 580Y mutations, though samples with wild-type and other non-synonymous *kelch13* mutations are also seen. A 17kb duplication is seen exclusively in samples from Pursat, all of which have 493H *kelch13* mutations. Similarly, a 80kb duplication is seen exclusively in Pailin in two samples which both have the 580Y *kelch13* mutation.

Amongst samples from ESEA carrying *kelch13* mutants, 223/555 (40%) have *plasmepsin 2-3* duplications, which is a significantly higher proportion than the 23/627 (3%) carrying wild-type *kelch13* (Fisher's exact $p=9.2x10^{-59}$). Similarly amongst samples from ESEA carrying *kelch13* mutants, 117/472 (24%) have *mdr1* duplications, which is a significantly higher proportion than the 19/559 (3%) carrying wild-type *kelch13* (Fisher's exact $p=3.2x10^{-25}$). However, of the 186 samples from ESEA carrying both *kelch13* mutations and *plasmepsin 2-3* duplications, only 12 (6%) also carry *mdr1* duplications, which is a significantly lower proportion than the 103/284 (36%) *kelch13* mutants without *plasmepsin 2-3* duplications (Fisher's exact $p=7.0x10^{-15}$), highlighting the previous reported antagonistic effect between *plasmepsin 2-3* and *mdr1* duplications[16].

#### No evidence of resistance to less commonly used antimalarials

Samples carrying the *dhfr* double mutant 16V and 108T have been associated with resistance to proguanil but this specific combination is completely absent other than in three samples from western Cambodia. Similarly, we do not see any evidence of resistance to atovaquone, resistance to which has been associated with mutations at amino acid 268 in mitochondrial gene *cytB*. This is compatible with the very limited and restricted usage of this drug so far, and with the notion that mutations in this gene are usually acquired within the course of the infection but are unlikely to be transmitted[17].

#### Supplementary references


1 Miotto O, Amato R, Ashley EA, *et al.* Genetic architecture of artemisinin-resistant Plasmodium falciparum. *Nat Genet* 2015; **47**: 226–34.

2 Wang Z, Cabrera M, Yang J, *et al.* Genome-wide association analysis identifies genetic loci associated with resistance to multiple antimalarials in Plasmodium falciparum from China-Myanmar border. *Sci Rep* 2016; **6**: 33891.

3 Takala-Harrison S, Clark TG, Jacob CG, *et al.* Genetic loci associated with delayed clearance of Plasmodium falciparum following artemisinin treatment in Southeast Asia. *Proc Natl Acad Sci U S A* 2013; **110**: 240–5.

4 Amambua-Ngwa A, Amenga-Etego L, Kamau E, *et al.* Major subpopulations of Plasmodium falciparum in sub-Saharan Africa. *Science (80- )* 2019; **365**: 813–6.

5 Tindall SM, Vallières C, Lakhani DH, Islahudin F, Ting K-N, Avery S V. Heterologous Expression of a Novel Drug Transporter from the Malaria Parasite Alters Resistance to Quinoline Antimalarials. *Sci Rep* 2018; **8**: 2464.

6 Kinga Modrzynska K, Creasey A, Loewe L, *et al.* Quantitative genome re-sequencing defines multiple mutations conferring chloroquine resistance in rodent malaria. *BMC Genomics* 2012; **13**: 106.

7 Cowell AN, Istvan ES, Lukens AK, *et al.* Mapping the malaria parasite druggable genome by using in vitro evolution and chemogenomics. *Science (80- )* 2018; **359**: 191–9.

8 Apinjoh TO, Mugri RN, Miotto O, *et al.* Molecular markers for artemisinin and partner drug resistance in natural Plasmodium falciparum populations following increased insecticide treated net coverage along the slope of mount Cameroon: Cross-sectional study. *Infect Dis Poverty* 2017; **6**: 136.

9 Chauvin P, Menard S, Iriart X, *et al.* Prevalence of *Plasmodium falciparum* parasites resistant to sulfadoxine/pyrimethamine in pregnant women in Yaoundé, Cameroon: emergence of highly resistant *pfdhfr / pfdhps* alleles. *J Antimicrob Chemother* 2015; **70**: 2566–71.

10 Oguike MC, Falade CO, Shu E, *et al.* Molecular determinants of sulfadoxine-pyrimethamine resistance in Plasmodium falciparum in Nigeria and the regional emergence of dhps 431V. *Int J Parasitol Drugs Drug Resist* 2016; **6**: 220–9.

11 Carvalho CMB, Ramocki MB, Pehlivan D, *et al.* Inverted genomic segments and complex triplication rearrangements are mediated by inverted repeats in the human genome. *Nat Genet* 2011; **43**: 1074–81.

12 Amato R, Lim P, Miotto O, *et al.* Genetic markers associated with dihydroartemisinin–piperaquine failure in Plasmodium falciparum malaria in Cambodia: a genotype–phenotype association study. *Lancet Infect Dis* 2017; **17**: 164–73.

13 Witkowski B, Duru V, Khim N, *et al.* A surrogate marker of piperaquine-resistant Plasmodium falciparum malaria: a phenotype-genotype association study. *Lancet Infect Dis* 2017; **17**: 174–83.

14 Imwong M, Suwannasin K, Kunasol C, *et al.* The spread of artemisinin-resistant Plasmodium falciparum in the Greater Mekong subregion: a molecular epidemiology observational study. *Lancet Infect Dis* 2017; **17**: 491–7.

15 Amato R, Pearson RD, Almagro-Garcia J, *et al.* Origins of the current outbreak of multidrug-resistant malaria in southeast Asia: a retrospective genetic study. *Lancet Infect Dis* 2018; **18**: 337–45.

16 Amaratunga C, Lim P, Suon S, *et al.* Dihydroartemisinin-piperaquine resistance in Plasmodium falciparum malaria in Cambodia: A multisite prospective cohort study. *Lancet Infect Dis* 2016; **16**: 357–65.

17 Goodman CD, Siregar JE, Mollard V, *et al.* Parasites resistant to the antimalarial atovaquone fail to transmit by mosquitoes. *Science (80- )* 2016; **352**: 349–53.

#### Supplementary tables

**Supplementary Table 1. Breakdown of analysis set samples by geography.** Sites are divided into eight regions as described in the main text. Note that samples from Mae Sot and Ranong in western Thailand have been assigned to the Western SE Asia (WSEA) region, whereas samples from Sisakhet in eastern Thailand have been assigned to the Eastern SE Asia (ESEA) region. 8 returning travellers were reported as returning/passed sample QC from Ghana (3/2), Kenya (2/1), Uganda (2/1) and Mozambique (1/1). 16 samples which were identified as lab strains were excluded from analysis.

| Region | Country | Site | Sequenced samples | Analysis set samples |
|---|---|---|---|---|
| *SAM* | **Colombia** | Buenaventura | 3 | 3 |
| | | Guapi | 4 | 4 |
| | | Quibdo | 3 | 3 |
| | | Tumaco | 6 | 6 |
| | **Peru** | Iquitos, Loreto Province | 11 | 11 |
| | | Loreto | 12 | 10 |
| *WAF* | **Benin** | Homel | 102 | 36 |
| | **Burkina Faso** | Bobo-Dioulasso | 57 | 56 |
| | **Cameroon** | Buea | 239 | 235 |
| | **Gambia** | Basse | 124 | 102 |
| | | Brikama | 123 | 116 |
| | | Madina Samako | 14 | 0 |
| | | Njaiyel | 16 | 1 |
| | **Ghana** | Cape-Coast | 101 | 100 |
| | | Kintampo | 61 | 44 |
| | | Navrongo | 841 | 705 |
| | **Guinea** | Faranah | 60 | 37 |
| | | Nzerekore | 137 | 112 |
| | **Ivory Coast** | Abobo | 31 | 31 |
| | | Koumassi | 19 | 19 |
| | | Yopougon | 20 | 20 |
| | **Mali** | Bamako | 164 | 162 |
| | | Bandiagara | 9 | 8 |
| | | Faladje | 173 | 157 |
| | | Kolle | 51 | 47 |
| | | Nioro du Sahel | 52 | 52 |
| | **Mauritania** | Aioun | 9 | 9 |
| | | Kobeni | 23 | 21 |
| | | Nema | 33 | 27 |
| | | Selibaby | 21 | 19 |
| | **Nigeria** | Bagdary | 34 | 24 |
| | | Ilorin | 8 | 5 |
| | **Senegal** | Pikine | 86 | 84 |
| *CAF* | **Congo DR** | Kinshasa | 366 | 344 |
| *EAF* | **Ethiopia** | Shewa Robit Town Health Centre | 15 | 10 |

| | | | | |
|---|---|---|---|---|
| | | West Arsi Zone | 19 | 11 |
| | **Kenya** | Kilifi | 68 | 49 |
| | | Kisumu | 34 | 34 |
| | | Kombewa | 27 | 26 |
| | **Madagascar** | Antsohihy | 6 | 5 |
| | | Farafangana | 1 | 1 |
| | | Maevatanana | 18 | 18 |
| | **Malawi** | Chikwawa | 300 | 221 |
| | | Zomba | 51 | 33 |
| | **Tanzania** | Mkuzi-Muheza | 160 | 152 |
| | | Morogoro | 34 | 32 |
| | | Muheza | 16 | 15 |
| | | Muleba | 61 | 52 |
| | | Nachingwea | 79 | 65 |
| | **Uganda** | Apac | 14 | 12 |
| ***SAS*** | **Bangladesh** | Bandarban | 42 | 28 |
| | | Ramu | 51 | 49 |
| ***WSEA*** | **Myanmar** | Bago | 94 | 61 |
| | | Kawthaung | 51 | 50 |
| | | Myitkyina | 28 | 26 |
| | | Pyin Oo Lwin | 23 | 22 |
| | | Thabeikkyin | 54 | 52 |
| | **Thailand** | Mae Sot | 935 | 848 |
| | | Ranong | 27 | 20 |
| ***ESEA*** | **Cambodia** | Pailin | 157 | 132 |
| | | Preah Vihear | 210 | 144 |
| | | Pursat | 539 | 376 |
| | | Ratanakiri | 243 | 194 |
| | | Tasanh | 65 | 50 |
| | **Laos** | Attapeu | 86 | 84 |
| | | Xepon | 45 | 36 |
| | **Thailand** | Sisakhet | 28 | 20 |
| | **Viet Nam** | Binh Phuoc | 126 | 114 |
| | | Phuoc Long | 138 | 112 |
| ***OCE*** | **Indonesia** | Timika | 92 | 80 |
| | **Papua New Guinea** | East Sepik | 53 | 48 |
| | | Madang | 56 | 44 |
| | | Milne Bay | 30 | 29 |
| ***Returning travellers*** | | *Various locations* | 8 | 5 |
| ***Lab samples*** | | *Various locations* | 16 | 0 |
| **Total** | | | **7,113** | **5,970** |

**Supplementary Table 2. Studies contributing samples.** Full details of the studies are available from the data release web page (http://www.malariagen.net/resource/26).

| Study ID | Study title | Contact | Samples | Sites |
|---|---|---|---|---|
| **1001-PF-ML-DJIMDE** | Developing the Community Project with partners in Mali | Abdoulaye Djimdé<br><br>Malaria Research and Training Centre, University of Science, Techniques and Technologies of Bamako, Mali | 96 | Bandiagara (Mali), Faladje (Mali), Kolle (Mali) |
| **1004-PF-BF-OUEDRAOGO** | Developing the Community Project with partners in Burkina Faso | Jean-Bosco Ouedraogo<br><br>Institut de Recherche en Sciences de la Santé, Burkina Faso | 57 | Bobo-Dioulasso (Burkina Faso) |
| **1006-PF-GM-CONWAY** | Genome-wide analysis of genetic variation in The Gambia | Alfred Amambua-Ngwa<br><br>Medical Research Council Unit, The Gambia | 79 | Brikama (Gambia) |
| **1007-PF-TZ-DUFFY** | Mother Offspring Malaria Study (MOMS) in Tanzania | Patrick Duffy<br><br>NIH National Institute of Allergy and Infectious Diseases (NIAID), USA | 50 | Morogoro (Tanzania), Muheza (Tanzania) |
| **1008-PF-SEA-RINGWALD** | Containment of artemisinin tolerant malaria parasites in South-East Asia (ARCE) | Pascal Ringwald<br><br>World Health Organization (WHO), Switzerland | 234 | Kawthaung (Myanmar), Phuoc Long (Viet Nam), Xepon (Laos) |

| | | | | |
|---|---|---|---|---|
| **1010-PF-TH-ANDERSON** | Genetic variation underlying drug resistance at the Thai-Burmese border | Tim J C Anderson<br><br>Texas Biomedical Research Institute, USA | 108 | Mae Sot (Thailand) |
| **1011-PF-KH-SU** | Genome-wide scans of cultured adapted parasites in Cambodia | Thomas E Wellems<br><br>NIH National Institute of Allergy and Infectious Diseases (NIAID), USA | 41 | Pursat (Cambodia) |
| **1012-PF-KH-WHITE** | Developing the Community Project with partners in Cambodia | Nicholas J White<br><br>Mahidol Oxford Tropical Medicine Research Unit (MORU), University of Oxford, Thailand | 2 | Pailin (Cambodia) |
| **1013-PF-PEGB-BRANCH** | Developing the Community Project with partners in Peru | Julian C Rayner<br><br>Wellcome Sanger Institute, UK | 16 | Iquitos, Loreto Province (Peru) |
| **1014-PF-SSA-SUTHERLAND** | Analysis of *Plasmodium falciparum* samples from UK travellers returning from malaria endemic countries | Colin Sutherland<br><br>London School of Hygiene and Tropical Medicine, UK | 8 | Ghana returning traveller (Ghana), Kenya returning traveller (Kenya), Mozambique returning traveller (Mozambique), Uganda returning traveller (Uganda) |
| **1015-PF-KE-NZILA** | Genome-wide association study of in vitro drug resistance in Kenya | Kevin Marsh<br><br>Nuffield Department of Medicine, University of Oxford, UK | 60 | Kilifi (Kenya) |
| **1016-PF-TH-NOSTEN** | Developing the Community Project with partners in Thailand | Francois Nosten<br> | 21 | Mae Sot (Thailand) |

| | | | | |
|---|---|---|---|---|
| | | Nuffield Department of Medicine, University of Oxford, UK | | |
| **1017-PF-GH-AMENGA-ETEGO** | Population genetics of natural populations in Northern Ghana | Lucas Amenga-Etego<br><br>Navrongo Health Research Centre, Ghana | 390 | Navrongo (Ghana) |
| **1020-PF-VN-BONI** | Measuring in vitro drug sensitivity in Vietnam | Tran Tinh Hien<br><br>Oxford University Clinical Research Unit (OUCRU), Vietnam | 24 | Binh Phuoc (Viet Nam) |
| **1021-PF-PG-MUELLER** | Building a national repository of malaria isolates in Papua New Guinea | Ivo Mueller<br><br>Barcelona Centre for International Health Research, Spain | 57 | East Sepik (Papua New Guinea), Madang (Papua New Guinea) |
| **1022-PF-MW-OCHOLLA** | Genome variation and selection in clinical isolates from rural Malawi | Alister Craig<br><br>Liverpool School of Tropical Medicine, UK | 351 | Chikwawa (Malawi), Zomba (Malawi) |
| **1023-PF-CO-ECHEVERRI-GARCIA** | Comparative analysis of permeome genes and drug resistance in Colombia | Diego F Echeverri-Garcia<br><br>International Center for Medical Research and Training - CIDEIM, Colombia | 17 | Buenaventura (Colombia), Guapi (Colombia), Quibdo (Colombia), Tumaco (Colombia) |
| **1024-PF-UG-BOUSEMA** | FightMal - Correlating protection from malaria with immune profile of infected individuals in Uganda | Teun Bousema<br><br>London School of Hygiene & Tropical Medicine, UK | 14 | Apac (Uganda) |
| **1026-PF-GN-CONWAY** | Effects of transmission intensity on population structure and signatures of selection in Guinea | David Conway<br> | 197 | Faranah (Guinea), Nzerekore (Guinea) |

| | | | | |
|---|---|---|---|---|
| | | London School of Hygiene & Tropical Medicine, UK | | |
| **1027-PF-KE-BULL** | Genomics of severe malaria and low host immunity in Kenya | Peter C Bull<br><br>Department of Pathology, University of Cambridge, UK | 11 | Kilifi (Kenya) |
| **1031-PF-SEA-PLOWE** | Artemisinin Resistance Confirmation, Characterization and Containment (ARC3) | Chris Plowe<br><br>Duke University, USA | 192 | Bandarban (Bangladesh), Mae Sot (Thailand), Pailin (Cambodia), Tasanh (Cambodia) |
| **1044-PF-KH-FAIRHURST** | Genomics of parasite clearance and recrudescence rates in Cambodia | Thomas E Wellems<br><br>NIH National Institute of Allergy and Infectious Diseases (NIAID), USA | 602 | Preah Vihear (Cambodia), Pursat (Cambodia), Ratanakiri (Cambodia) |
| **1052-PF-TRAC-WHITE** | Tracking Resistance to Artemisinin Collaboration (TRAC) | Elizabeth Ashley<br><br>Mahidol Oxford Tropical Medicine Research Unit (MORU), University of Oxford, Thailand | 1,172 | Attapeu (Laos), Bago (Myanmar), Binh Phuoc (Viet Nam), Ilorin (Nigeria), Kinshasa (Congo DR), Mae Sot (Thailand), Myitkyina (Myanmar), Pailin (Cambodia), Preah Vihear (Cambodia), Pursat (Cambodia), Pyin Oo Lwin (Myanmar), Ramu (Bangladesh), Ranong (Thailand), Ratanakiri (Cambodia), Sisakhet (Thailand), Thabeikkyin (Myanmar) |
| **1062-PF-PG-BARRY** | Understanding malaria parasite populations and outbreaks in Papua New Guinea | Alyssa Barry<br> | 82 | East Sepik (Papua New Guinea), Milne Bay (Papua New Guinea) |

| | | | | |
|---|---|---|---|---|
| | | Walter and Eliza Hall Institute, Australia | | |
| **1083-PF-GH-CONWAY** | Alternative molecular mechanisms for erythrocyte invasion by *P. falciparum* in Ghana | Gordon Awandare<br><br>University of Ghana, Legon, Ghana | 101 | Kintampo (Ghana), Navrongo (Ghana) |
| **1093-PF-CM-APINJOH** | Population genetics of *P. falciparum* parasites in South-Western Cameroon | Tobias Apinjoh<br><br>University of Buea, Cameroon | 239 | Buea (Cameroon) |
| **1094-PF-GH-AMENGA-ETEGO** | Population genetics of *P. falciparum* parasites in Northern Ghana | Lucas Amenga-Etego<br><br>Navrongo Health Research Centre, Ghana | 256 | Navrongo (Ghana) |
| **1095-PF-TZ-ISHENGOMA** | Genome variation and its effect on ACT treatment outcome in Tanzania | Deus Ishengoma<br><br>National Institute for Medical Research (NIMR), United Republic of Tanzania | 300 | Mkuzi-Muheza (Tanzania), Muleba (Tanzania), Nachingwea (Tanzania) |
| **1096-PF-GH-GHANSAH** | Population genetics of *P. falciparum* parasites in Southern Ghana | Anita Ghansah<br><br>Nogouchi Memorial Institute for Medical Research, Legon-Accra, Ghana | 101 | Cape-Coast (Ghana) |
| **1097-PF-ML-MAIGA** | Detection of artemisinin-resistant *Plasmodium falciparum* parasites in Southern Mali | Abdoulaye Djimdé<br><br>Malaria Research and Training Centre, University of Science, Techniques and Technologies of Bamako, Mali | 137 | Faladje (Mali) |

| | | | | |
|---|---|---|---|---|
| **1098-PF-ET-GOLASSA** | The prevalence of asymptomatic carriage; emergence of parasite mutations conferring anti-malaria drug resistance; and G6PD deficiency in the human population, as possible impediments to malaria elimination in Ethiopia | Lemu Golassa<br><br>Aklilu Lemma Institute of Pathobiology, Addis Ababa University, Ethiopia | 34 | Shewa Robit Town Health Centre (Ethiopia), West Arsi Zone (Ethiopia) |
| **1100-PF-CI-YAVO** | Drug resistance and *Plasmodium falciparum* diversity in forest zone of Côte d'Ivoire | William Yavo<br><br>Malaria Research and Control Center of the National Institute of Public Health, Côte d'Ivoire | 70 | Abobo (Ivory Coast), Koumassi (Ivory Coast), Yopougon (Ivory Coast) |
| **1101-PF-CD-ONYAMBOKO** | Efficacy of 3 ACTs in treating *falciparum* malaria in the Democratic Republic of Congo | Caterina A. Fanello<br><br>Mahidol Oxford Tropical Medicine Research Unit (MORU), University of Oxford, Thailand | 174 | Kinshasa (Congo DR) |
| **1102-PF-MG-RANDRIANARIVELOJOSIA** | Genotyping *P. falciparum* and *P. vivax* in Madagascar | Milijaona Randrianarivelojosia<br><br>Institut Pasteur de Madagascar | 25 | Antsohihy (Madagascar), Farafangana (Madagascar), Maevatanana (Madagascar) |
| **1103-PF-PDN-GMSN-NGWA** | Population genetics of cross-border *P. falciparum* parasites in West Africa | Alfred Amambua-Ngwa<br><br>Medical Research Council Unit, The Gambia | 34 | Bagdary (Nigeria) |
| **1107-PF-KEN-KAMAU** | Population genetics of *P. falciparum* parasites in Kenya | Edwin Kamau<br><br>Department of Pathology and Laboratory Services, Walter Reed National Military Medical Center, Bethesda, MD USA | 61 | Kisumu (Kenya), Kombewa (Kenya) |
| **1125-PF-TH-NOSTEN** | Investigating artemisinin resistance emergence on Thai-Burmese border | Francois Nosten | 674 | Mae Sot (Thailand) |

| | | | | |
|---|---|---|---|---|
| | |<br>Nuffield Department of Medicine, University of Oxford, UK | | |
| **1127-PF-ML-SOULEYMANE** | Genetic analysis of *P. falciparum* before and after artemether-lumefantrine treatment in Mali | Abdoulaye Djimdé<br><br>Malaria Research and Training Centre, University of Science, Techniques and Technologies of Bamako, Mali | 164 | Bamako (Mali) |
| **1131-PF-BJ-BERTIN** | Identification of virulence factors in cerebral malaria in Benin | Gwladys Bertin<br><br>Research institute for development, France | 102 | Homel (Benin) |
| **1134-PF-ML-CONWAY** | Population Genetics of *P. falciparum* in West Africa | David Conway<br><br>London School of Hygiene & Tropical Medicine, UK | 52 | Nioro du Sahel (Mali) |
| **1135-PF-SN-CONWAY** | Parasite adaption in Senegal at molecular, functional and population level | David Conway<br><br>London School of Hygiene & Tropical Medicine, UK | 86 | Pikine (Senegal) |
| **1136-PF-GM-NGWA** | *Plasmodium falciparum* anti-malarial drug resistance in the Gambia: Identification of potential genetic markers by retrospective whole genome approaches | Alfred Amambua-Ngwa<br><br>Medical Research Council Unit, The Gambia | 100 | Basse (Gambia), Brikama (Gambia) |
| **1137-PF-GM-DALESSANDRO** | Malaria transmission dynamics in The Gambia: Defining the spatial and temporal spread of malaria at micro-level (village) | Alfred Amambua-Ngwa<br><br>Medical Research Council Unit, The Gambia | 68 | Basse (Gambia) |

| | | | | |
|---|---|---|---|---|
| **1138-PF-CD-FANELLO** | Parenteral artesunate compared to quinine as a cause of late post-treatment anaemia in African children with *hyperparasitaemic P. falciparum* malaria (DHART) | Caterina A. Fanello<br><br>Mahidol Oxford Tropical Medicine Research Unit (MORU), University of Oxford, Thailand | 77 | Kinshasa (Congo DR) |
| **1141-PF-GM-CLAESSENS** | Genomic characterization of *P. falciparum* from asymptomatic infections in The Gambia | Antoine Claessens<br><br>Medical Research Council Unit, The Gambia | 31 | Madina Samako (Gambia), Njaiyel (Gambia) |
| **1145-PF-PE-GAMBOA** | Genotype-phenotype study of erythrocyte invasion in Peruvian *P. falciparum* isolates | Dionicia Gamboa<br><br>Universidad Peruana Cayetano Heredia (UPCH), Peru | 13 | Loreto (Peru) |
| **1146-PF-MULTI-PRICE** | Characterisation of drug resistance in Indonesian *P. falciparum* populations | Sarah Auburn<br><br>Menzies School of Health Research, Australia | 92 | Timika (Indonesia) |
| **1147-PF-MR-CONWAY** | Population genetics of *P. falciparum* parasites in Mauritania | David Conway<br><br>London School of Hygiene & Tropical Medicine, UK | 86 | Aioun (Mauritania), Kobeni (Mauritania), Nema (Mauritania), Selibaby (Mauritania) |
| **1151-PF-GH-AMENGA-ETEGO** | Testing the effectiveness of selective whole genome amplification on samples collected in Northern Ghana | Lucas Amenga-Etego<br><br>Navrongo Health Research Centre, Ghana | 155 | Navrongo (Ghana) |
| **Total** | | | **7,113** | |

**Supplementary Table 3. Summary of discovered variant positions.** We divide variant positions into those containing single nucleotide polymorphisms (SNPs) and non-SNPs (indels and combinations of SNPs and indels at the same position). We then further sub-divide each of these into those within exons (coding) and those in intronic or intergenic regions (non-coding). We further sub-divide SNPs into those containing only two alleles (bi-allelic) or those contains three or more alleles (multi-allelic). Discovered variant positions are unique positions in the reference genome where either SNP or indel variation was discovered by our analysis pipeline. Pass variant positions are the subset of discovered positions that passed our quality filters. Alleles per pass position shows the mean number of distinct alleles at each pass position; biallelic variants have 2 alleles have two alleles by definition.

| Type | Coding | Multi-allelic | Discovered variant positions | Pass variant positions | % pass | Alleles per pass position |
|---|---|---|---|---|---|---|
| SNP | Coding | Bi-allelic | 1,590,717 | 1,042,291 | 66% | 2.0 |
| | | Multi-allelic | 195,356 | 139,388 | 71% | 3.1 |
| | Non-coding | Bi-allelic | 1,203,255 | 581,976 | 48% | 2.0 |
| | | Multi-allelic | 179,393 | 72,064 | 40% | 3.1 |
| non-SNP | Coding | | 882,235 | 326,199 | 37% | 3.6 |
| | Non-coding | | 2,000,740 | 949,072 | 47% | 3.5 |
| Total | | | 6,051,696 | 3,110,990 | 51% | 2.7 |

**Supplementary Table 4. Breakpoints of duplications of *gch1*.** Breakpoint IDs are shown in the first column and can be used to match to the per sample breakpoints in the data release. Breakpoints are generally poly-A or poly-T repeats and Breakpoint 1 and Breakpoint 2 show the start and end of each repeat in the reference genome. Breakpoints 3 and 4 are shown for DUP-TRPINV-DUP breakpoint. The Sites column shows the sites where this breakpoint was identified, together with the number of QC pass samples at that site that had this set of breakpoints. DHFR/DHPS shows amino acid haplotypes at the amino acids *dhfr* (51, 59 and 108) and *dhps* (437 and 540) seen in samples with each breakpoint, together with the number of samples for each haplotype in brackets. Note we only show these haplotypes for samples that were homozygous for the haplotype.

| Breakpoint ID | Breakpoint 1 | Breakpoint 2 | Breakpoint 3 | Breakpoint 4 | Sites | DHFR/DHPS |
|---|---|---|---|---|---|---|
| **PfGCH1_dup_1** | 968847-968881 | 977926-977949 | | | Kinshasa (6), Apac (1), Binh Phuoc (2), Pailin (9), Pursat (39), Ratanakiri (2), Tasanh (1), Bandarban (2), Ramu (1), Abobo (1), Buea (6), Kintampo (1), Navrongo (2), Bago (9), Kawthaung (19), Mae Sot (277), Myitkyina (3), Ranong (1), Thabeikkyin (1) | IRN/AK (2), IRN/GE (280), IRN/GK (20), IRN/GN (24), NRN/GE (19), NRN/GK (3), NRN/GN (1) |
| **PfGCH1_dup_2** | 946265-946284 | 980622-980659 | | | Pailin (3) | NCT/AK (3) |
| **PfGCH1_dup_3** | 959516-959540 | 978164-978191 | | | Pursat (5), Cape-Coast (1) | IRN/GK (5), IRN/GN (1) |
| **PfGCH1_dup_4** | 970992-971023 | 975712-975747 | | | Kintampo (1), Navrongo (1) | IRN/AK (1) |
| **PfGCH1_dup_5** | 953141-953174 | 978164-978191 | | | West Arsi Zone (2) | IRN/GE (2) |
| **PfGCH1_dup_6** | 959516-959540 | 981032-981060 | | | Brikama (1), Buea (1) | IRN/AK (1) |
| **PfGCH1_dup_7** | 974100-974119 | 986443-986465 | | | Maevatanana (3) | IRN/GE (1), IRN/GK (2) |

| | | | | | | |
|---|---|---|---|---|---|---|
| **PfGCH1_dup_8** | 973800-973825 | 976004-976045 | | | Attapeu (2), Binh Phuoc (8), Pailin (13), Phuoc Long (11), Preah Vihear (3), Pursat (4), Ratanakiri (5), Sisakhet (2), Tasanh (2), Ramu (1), Abobo (4), Brikama (2), Buea (21), Cape-Coast (3), Homel (1), Kintampo (1), Koumassi (2), Navrongo (11), Nioro du Sahel (1), Bago (7), Kawthaung (6), Mae Sot (22), Pyin Oo Lwin (2), Ranong (4), Thabeikkyin (1) | IRN/AK (1), IRN/GE (50), IRN/GK (39), IRN/GN (29), NCS/AK (1), NCS/GK (3), NRN/GE (2), NRN/GK (3) |
| **PfGCH1_dup_9** | 968847-968881 | 976155-976170 | | | Bandarban (2), Ramu (5), Navrongo (9), Bago (4), Kawthaung (14), Mae Sot (73), Myitkyina (4), Pyin Oo Lwin (10), Ranong (2), Thabeikkyin (2) | IRN/GE (86), IRN/GK (5), IRN/GN (4), NRN/AK (2), NRN/GE (18), NRN/GK (1) |
| **PfGCH1_DTD_1** | 929743-929759 | 940895-940912 | 978164-978191 | 980075-980103 | Guapi (1) | NCN/AK (1) |
| **PfGCH1_DTD_2** | 938785-938805 | 968847-968881 | 977926-977949 | 980363-980386 | Milne Bay (4), Timika (32), Mae Sot (1) | IRNGE (1), NRNGE (35) |

**Supplementary Table 5. Breakpoints of duplications of *mdr1*.** Breakpoint IDs are shown in the first column and can be used to match to the per sample breakpoints in the data release. Breakpoints are generally poly-A or poly-T repeats and Breakpoint 1 and Breakpoint 2 show the start and end of each repeat in the reference genome. Breakpoints 3 and 4 are shown for DUP-TRPINV-DUP breakpoints. The Sites column shows the sites where this breakpoint was identified, together with the number of QC pass samples at that site that had this set of breakpoints. K13 shows mutations in the *kelch13* gene seen in samples with each breakpoint, together with the number of samples for each mutation in brackets. Note we only show these mutations for samples that were homozygous for the mutation. WT=wild-type.

| Breakpoint ID | Breakpoint 1 | Breakpoint 2 | Breakpoint 3 | Breakpoint 4 | Sites | K13 |
|---|---|---|---|---|---|---|
| **PfMDR1_dup_1** | 938329-938357 | 980012-980040 | | | Kawthaung (4) | C580Y (4) |
| **PfMDR1_dup_2** | 949514-949527 | 967253-967266 | | | Mae Sot (9) | WT (5), M476I (1), P441L (3) |
| **PfMDR1_dup_3** | 947790-947800 | 962444-962454 | | | Mae Sot (67) | WT (33), A675V (2), C580Y (12), G538V (10), N458Y (7), P527H (1) |
| **PfMDR1_dup_4** | 953961-953982 | 973008-973034 | | | Bago (1), Mae Sot (63) | WT (47), A481V (1), C580Y (14) |
| **PfMDR1_dup_5** | 953961-953982 | 965409-865428 | | | Mae Sot (21) | WT (11), C580Y (4), P574L (3) |
| **PfMDR1_dup_6** | 947968-947986 | 969783-969812 | | | Pailin (5), Phuoc Long (2), Preah Vihear (3), Pursat (17), Sisakhet (10), Tasanh (4), Bago (3), Mae Sot (2), Ranong (1) | WT (5), C580Y (10), R539T (28), Y493H (1), c580y (1) |

| | | | | | | |
|---|---|---|---|---|---|---|
| **PfMDR1_dup_7** | 953961-953982 | 970215-970252 | | | Pailin (1), Phuoc Long (2), Preah Vihear (4), Pursat (11), Tasanh (1), Kawthaung (2), Mae Sot (98) | WT (53), A675V (5), C580Y (23), K479I (1), N458Y (2), P441L (4), P443S (1), P527H (1), P553L (1), R539T (3), R561H (6), Y493H (9) |
| **PfMDR1_dup_8** | 780906-780927 | 980012-980040 | | | | |
| **PfMDR1_dup_9** | 870473-870502 | 964628-964646 | | | Mae Sot (1) | |
| **PfMDR1_dup_10** | 795494-795527 | 964505-964540 | | | Kawthaung (1) | WT (1) |
| **PfMDR1_dup_11** | 888324-888349 | 970215-970252 | | | | |
| **PfMDR1_dup_12** | 868667-868699 | 964505-964540 | | | Pailin (3) | WT (2) |
| **PfMDR1_dup_13** | 946696-946717 | 964505-964540 | | | Binh Phuoc (4), Pailin (9), Phuoc Long (3), Preah Vihear (6), Pursat (28), Ratanakiri (1), Sisakhet (1), Bago (2), Mae Sot (1) | WT (10), C580Y (26), R539T (6), Y493H (6) |
| **PfMDR1_dup_14** | 946696-946717 | 970215-970252 | | | | |
| **PfMDR1_dup_15** | 946346-946375 | 970215-970252 | | | Pailin (1), Tasanh (1), Milne Bay (2), Kawthaung (1), Mae Sot (5), Ranong (2) | WT (6), C580Y (2), K479I (1), P574L (1), R539T (1) |
| **PfMDR1_dup_16** | 948197-948217 | 976430-976456 | | | Mae Sot (8) | WT (8) |
| **PfMDR1_dup_17** | 946346-946375 | 986036-986067 | | | Kawthaung (1) | WT (1) |
| **PfMDR1_dup_18** | 953961-953982 | 976144-976170 | | | Pursat (1), Mae Sot (2) | WT (2), C580Y (1) |
| **PfMDR1_dup_19** | 953961-953982 | 989700-989721 | | | | |
| **PfMDR1_dup_20** | 946696-946717 | 969783-969812 | | | Pursat (1) | C580Y (1) |

| | | | | | | |
|---|---|---|---|---|---|---|
| **PfMDR1_dup_21** | 946346-946375 | 973140-973162 | | | Pursat (5) | WT (5) |
| **PfMDR1_dup_22** | 943418-943437 | 970215-970252 | | | Mae Sot (1) | A675V (1) |
| **PfMDR1_dup_23** | 942351-942376 | 970215-970252 | | | Mae Sot (1) | C580Y (1) |
| **PfMDR1_dup_24** | 939060-939083 | 976144-976170 | | | Bago (1) | WT (1) |
| **PfMDR1_dup_25** | 938329-938357 | 973140-973162 | | | Thabeikkyin (1) | |
| **PfMDR1_dup_26** | 953961-953982 | 976430-976456 | | | Mae Sot (1), Myitkyina (1) | WT (1), F446I (1) |
| **PfMDR1_dup_27** | 937002-937024 | 969783-969812 | | | Pyin Oo Lwin (7) | P574L (7) |
| **PfMDR1_DTD_1** | 928340-928359 | 938911-938930 | 964505-964532 | 985396-985423 | Pursat (4) | Y493H (4) |

**Supplementary Table 6. Breakpoints of duplications of *plasmepsin 2-3*.** Breakpoint IDs are shown in the first column and can be used to match to the per sample breakpoints in the data release. Breakpoints are generally poly-A or poly-T repeats and Breakpoint 1 and Breakpoint 2 show the start and end of each repeat in the reference genome. The Sites column shows the sites where this breakpoint was identified, together with the number of QC pass samples at that site that had this set of breakpoints. K13 shows mutations in the *kelch13* gene seen in samples with each breakpoint, together with the number of samples for each mutation in brackets. Note we only show these mutations for samples that were homozygous for the mutation. WT=wild-type.

| Breakpoint ID | Breakpoint 1 | Breakpoint 2 | Sites | K13 |
|---|---|---|---|---|
| **PfPlasmepsin_1** | 289611-289621 | 298782-298792 | Pailin (49), Preah Vihear (19), Pursat (173), Ratanakiri (3), Sisakhet (1), Tasanh (11), Mae Sot (3) | WT (26), C580Y (208), F395Y (1), H719N (2), Y493H (6) |
| **PfPlasmepsin_2** | 283034-283069 | 300493-300522 | Pursat (4) | Y493H (4) |
| **PfPlasmepsin_3** | 283034-283069 | 362990-363020 | Pailin (2) | C580Y (2) |

**Supplementary Table 7. Genes ranked by global differentiation score.** The table contains the ten genes with highest global differentiation score. The full list of all genes is available in the data release (https://www.malariagen.net/resource/26). Gene=GeneDB ID. Name=GeneDB name. Mut=highest $F_{ST}$ non-synonymous SNP within the gene. $F_{ST}$ = $F_{ST}$ of mutation in Mut column. NRAF=non-reference allele frequency in each region of the mutation shown in the Mut column. SAM=South America, WAF=West Africa, CAF=Central Africa, EAF=East Africa, SAS=South Asia, WSEA=West south-east Asia, ESEA=East south-east Asia, OCE=Oceania. Score=global differentiation score (see Methods).

| | | | | NRAF | | | | | | | | |
|---|---|---|---|---|---|---|---|---|---|---|---|---|
| **Gene** | **Name** | **Mut** | **$F_{ST}$** | **SAM** | **WAF** | **CAF** | **EAF** | **SAS** | **WSEA** | **ESEA** | **OCE** | **Score** |
| **PF3D7_1346800** | P47 | S242L and V247A | 1.000 | 1.00 | 0.00 | 0.00 | 0.00 | 0.00 | 0.00 | 0.00 | 0.00 | 1 |
| **PF3D7_0207600** | SERA5 | K383N | 1.000 | 1.00 | 0.00 | 0.00 | 0.00 | 0.00 | 0.00 | 0.00 | 0.00 | 0.95 |
| **PF3D7_0935600** | GIG | G171D | 0.993 | 0.00 | 0.00 | 0.00 | 0.00 | 0.95 | 1.00 | 1.00 | 0.97 | 0.90 |
| **PF3D7_0406200** | Pfs16 | S90N | 0.990 | 0.00 | 0.00 | 0.00 | 0.00 | 0.93 | 0.99 | 1.00 | 1.00 | 0.88 |
| **PF3D7_0315200** | CTRP | D319N | 0.989 | 0.97 | 0.00 | 0.00 | 0.00 | 0.99 | 0.99 | 1.00 | 0.98 | 0.87 |
| **PF3D7_1361100** | SEC24A | S301P | 0.989 | 0.16 | 0.00 | 0.00 | 0.00 | 0.95 | 1.00 | 0.99 | 0.98 | 0.86 |
| **PF3D7_1116800** | HSP101 | R172S | 0.989 | 0.22 | 0.00 | 0.00 | 0.01 | 0.92 | 1.00 | 1.00 | 1.00 | 0.85 |
| **PF3D7_0320400** | Cap380 | W2127R | 0.986 | 0.00 | 0.00 | 0.00 | 0.00 | 0.00 | 0.00 | 0.00 | 0.98 | 0.83 |
| **PF3D7_0709000** | CRT | C72S | 0.980 | 0.00 | 0.00 | 0.00 | 0.00 | 0.00 | 0.00 | 0.00 | 0.98 | 0.82 |
| **PF3D7_1344300** | | E497Q | 0.980 | 0.19 | 0.00 | 0.00 | 0.00 | 0.90 | 1.00 | 0.99 | 0.93 | 0.81 |

**Supplementary Table 8. Genes ranked by local differentiation score.** The table contains the ten genes with highest local differentiation score. We have excluded genes that are within 50kb of a gene with a higher local differentiation score. The full list of all genes is available in the data release (https://www.malariagen.net/resource/26). Gene=GeneDB ID. Name=GeneDB name. Columns WAF to OCE show the most highly differentiated non-synonymous SNP in each region. Numbers in brackets show rank amongst all non-synonymous SNPs. A hyphen indicates that there were no segregating SNPs in the gene in this region. SAM=South America, WAF=West Africa, CAF=Central Africa, EAF=East Africa, SAS=South Asia, WSEA=West south-east Asia, ESEA=East south-east Asia, OCE=Oceania. Score=local differentiation score (see Methods).

| Gene | Name | WAF | EAF | SAS | WSEA | ESEA | OCE | Score |
|---|---|---|---|---|---|---|---|---|
| **PF3D7_1318100** | . | P28L (38273) | D193Y (25047) | E125D (2406) | D193Y (129) | D193Y (1) | D193Y (3) | 0.91 |
| **PF3D7_0709000** | CRT | I356T (7) | Q271E (56) | I356T (82) | N326S (1226) | N326S (2) | T333S (1097) | 0.85 |
| **PF3D7_1012900** | ATG18 | K301N (25704) | K335Q (28852) | - | T38I (291) | T38I (7) | T38I (1) | 0.85 |
| **PF3D7_0523000** | MDR1 | S1082A (305) | N86Y (27) | S784L (3068) | F1226Y (995) | Y184F (8) | N1042D (2) | 0.84 |
| **PF3D7_0525100** | ACS10 | N341I (9) | M300I (5) | D170N (1557) | G263S (534) | T172I (807) | P127Q (2181) | 0.83 |
| **PF3D7_0810800** | PPPK-DHPS | I431V (12) | A581G (4) | G437A (1705) | A581G (154) | A581G (132) | G437A (251) | 0.81 |
| **PF3D7_1218400** | . | E12D (2613) | R294C (5889) | R294C (18) | R294C (1665) | R294C (23) | - | 0.75 |
| **PF3D7_0404500** | P52 | Q69E (26) | N352K (536) | I473L (1150) | T416I (6) | I473L (2901) | M87I (344) | 0.74 |
| **PF3D7_1222400** | ApiAP2 | C1111S (160) | V740I (12) | E26K and V132G (1252) | T2125N (3999) | Q1489H (26) | H662N (97) | 0.74 |
| **PF3D7_1343700** | K13 | D109H (6417) | K189T (18110) | K189T (2890) | F446I (4) | C580Y (28) | K189T (2059) | 0.74 |

**Supplementary Table 9. Number of samples used to determine proportions in Table 2.**

| Marker | Associated with resistance to | South America | West Africa | Central Africa | East Africa | South Asia | West south-east Asia | East south-east Asia | Oceania |
|---|---|---|---|---|---|---|---|---|---|
| ***crt***<br>76T | Chloroquine | 37 | 1,910 | 262 | 697 | 73 | 1,075 | 1,245 | 195 |
| ***dhfr***<br>108N | Pyrimethamine | 37 | 1,943 | 342 | 733 | 76 | 1,079 | 1,256 | 200 |
| ***dhps***<br>437G | Sulfadoxine | 37 | 1,901 | 331 | 702 | 62 | 1,078 | 1,201 | 197 |
| ***mdr1***<br>2+ copies | Mefloquine | 33 | 2,050 | 309 | 678 | 63 | 950 | 1,055 | 185 |
| ***kelch13***<br>WHO list | Artemisinin | 37 | 2,198 | 335 | 732 | 77 | 1,027 | 1,195 | 199 |
| ***plasmepsin 2-3***<br>2+ copies | Piperaquine | 36 | 2,216 | 342 | 736 | 76 | 1,076 | 1,176 | 201 |
| ***dhfr***<br>triple mutant | SP (treatment) | 37 | 1,851 | 283 | 693 | 65 | 1,042 | 1,221 | 201 |
| ***dhfr* and *dhps***<br>sextuple mutant | SP (IPTp) | 37 | 2,228 | 338 | 701 | 68 | 906 | 867 | 201 |
| ***kelch13* and *mdr1*** | AS-MQ | 37 | 2,230 | 343 | 738 | 77 | 1,013 | 1,128 | 201 |
| ***kelch13* and *plasmepsin 2-3*** | DHA-PPQ | 37 | 2,231 | 344 | 739 | 77 | 1,078 | 1,188 | 201 |

**Supplementary Table 10. Frequencies of mutations associated with mono- and multi-drug resistance pre- and post-2011.** The first column shows the gene and marker used to detect resistance. The second column shows the drug the markers are associated with resistance to. The remaining columns show the proportion of samples within each region that were associated with resistance to each drug. The upper number shows samples collected between 2001-2011 and the lower number samples collected between 2012-2015. The number of samples (n) used to create each proportion varies by drug due to differential missingness among markers. This table includes all samples from the date range 2001-2015, though note that prior to 2007 we had only sequenced samples from Western SE Asia. A hyphen indicates that no samples from the region were available in the date range.

| Marker | Associated with resistance to | South America | West Africa | Central Africa | East Africa | South Asia | West Southeast Asia | East Southeast Asia | Oceania |
|---|---|---|---|---|---|---|---|---|---|
| ***crt***<br>76T | Chloroquine | 1.00 (n=37)<br>- | 0.43 (n=621)<br>0.40 (n=1289) | -<br>0.66 (n=262) | 0.12 (n=356)<br>0.17 (n=341) | 0.88 (n=26)<br>0.96 (n=47) | 1.00 (n=717)<br>0.99 (n=358) | 0.96 (n=893)<br>0.99 (n=352) | 0.98 (n=63)<br>0.99 (n=132) |
| ***dhfr***<br>108N | Pyrimethamine | 0.97 (n=37)<br>- | 0.80 (n=609)<br>0.87 (n=1334) | -<br>1.00 (n=342) | 0.99 (n=361)<br>0.98 (n=372) | 1.00 (n=27)<br>1.00 (n=49) | 1.00 (n=721)<br>1.00 (n=358) | 0.99 (n=901)<br>1.00 (n=355) | 0.98 (n=66)<br>1.00 (n=134) |
| ***dhps***<br>437G | Sulfadoxine | 0.30 (n=37)<br>- | 0.78 (n=587)<br>0.74 (n=1314) | -<br>0.97 (n=331) | 0.95 (n=352)<br>0.91 (n=350) | 0.90 (n=21)<br>1.00 (n=41) | 1.00 (n=720)<br>1.00 (n=358) | 0.86 (n=861)<br>0.91 (n=340) | 0.45 (n=66)<br>0.69 (n=131) |
| ***mdr1***<br>2+ copies | Mefloquine | 0.00 (n=33)<br>- | 0.00 (n=611)<br>0.00 (n=1439) | -<br>0.00 (n=309) | 0.00 (n=304)<br>0.00 (n=374) | 0.00 (n=16)<br>0.00 (n=47) | 0.45 (n=635)<br>0.42 (n=315) | 0.15 (n=760)<br>0.05 (n=295) | 0.00 (n=58)<br>0.02 (n=127) |
| ***kelch13***<br>WHO list | Artemisinin | 0.00 (n=37)<br>- | 0.00 (n=723)<br>0.00 (n=1475) | -<br>0.00 (n=335) | 0.00 (n=361)<br>0.00 (n=371) | 0.00 (n=28)<br>0.00 (n=49) | 0.12 (n=696)<br>0.60 (n=331) | 0.45 (n=858)<br>0.50 (n=337) | 0.00 (n=67)<br>0.00 (n=132) |
| ***plasmepsin 2-3***<br>2+ copies | Piperaquine | 0.00 (n=36)<br>- | 0.00 (n=728)<br>0.00 (n=1488) | -<br>0.00 (n=342) | 0.00 (n=362)<br>0.00 (n=374) | 0.00 (n=27)<br>0.00 (n=49) | 0.00 (n=718)<br>0.00 (n=358) | 0.11 (n=863)<br>0.34 (n=313) | 0.00 (n=67)<br>0.00 (n=134) |
| ***dhfr***<br>triple mutant | SP (treatment) | 0.00 (n=37)<br>- | 0.65 (n=570)<br>0.79 (n=1281) | -<br>0.82 (n=283) | 0.93 (n=345)<br>0.90 (n=348) | 0.33 (n=24)<br>0.49 (n=41) | 0.91 (n=690)<br>0.88 (n=352) | 0.91 (n=875)<br>0.94 (n=346) | 0.00 (n=67)<br>0.00 (n=134) |
| ***dhfr* and *dhps***<br>sextuple mutant | SP (IPTp) | 0.00 (n=37)<br>- | 0.00 (n=735)<br>0.00 (n=1493) | -<br>0.01 (n=338) | 0.02 (n=354)<br>0.18 (n=347) | 0.15 (n=26)<br>0.21 (n=42) | 0.86 (n=576)<br>0.77 (n=330) | 0.19 (n=640)<br>0.21 (n=227) | 0.00 (n=67)<br>0.00 (n=134) |
| ***kelch13* and *mdr1*** | AS-MQ | 0.00 (n=37)<br>- | 0.00 (n=736)<br>0.00 (n=1494) | -<br>0.00 (n=343) | 0.00 (n=364)<br>0.00 (n=374) | 0.00 (n=28)<br>0.00 (n=49) | 0.04 (n=696)<br>0.33 (n=317) | 0.11 (n=810)<br>0.03 (n=318) | 0.00 (n=67)<br>0.00 (n=134) |
| ***kelch13* and p*lasmepsin* 2-3** | DHA-PPQ | 0.00 (n=37)<br>- | 0.00 (n=737)<br>0.00 (n=1494) | -<br>0.00 (n=344) | 0.00 (n=365)<br>0.00 (n=374) | 0.00 (n=28)<br>0.00 (n=49) | 0.00 (n=720)<br>0.00 (n=358) | 0.09 (n=869)<br>0.30 (n=319) | 0.00 (n=67)<br>0.00 (n=134) |

**Supplementary Table 11. Frequency of *crt* amino acid 72-76 haplotypes.** Here we have only included samples for which we have a homozygous call at amino acid 76, i.e. for which we could assign a chloroquine resistance phenotype. Mutant amino acids are shown in **underlined bold** font, wild-type in normal font. The first row shows the wild-type CVMNK haplotype. This is the only haplotype that does not have the 76T mutation and as such is the only haplotype considered sensitive to chloroquine. Rows 2-7 show other haplotypes that are seen as homozygotes. We have considered these resistant to chloroquine. Other samples are either heterozygous between different mutant haplotypes, or else the full 5-amino acid haplotype could not be resolved although the sample had the 76T mutation. These samples are included in the '*Other*' row and are also considered resistant to chloroquine. The final row shows the sum across rows 2-8 and corresponds to the first row of Table 2. SAM=South America, WAF=West Africa, CAF=Central Africa, EAF=East Africa, SAS=South Asia, WSEA=West south-east Asia, ESEA=East south-east Asia, OCE=Oceania.

| ***crt*** **72-76 haplotype** | **SAM (n=37)** | **WAF (n=1910)** | **CAF (n=262)** | **EAF (n=697)** | **SAS (n=73)** | **WSEA (n=1075)** | **ESEA (n=1245)** | **OCE (n=195)** |
|---|---|---|---|---|---|---|---|---|
| CVMNK | 0.00 | 0.59 | 0.34 | 0.86 | 0.07 | 0.00 | 0.03 | 0.01 |
| CVMN**T** | 0.27 | 0.00 | 0.00 | 0.00 | 0.00 | 0.00 | 0.00 | 0.00 |
| **S**VMN**T** | 0.30 | 0.00 | 0.00 | 0.00 | 0.00 | 0.00 | 0.00 | 0.99 |
| CVM**ET** | 0.43 | 0.00 | 0.00 | 0.00 | 0.00 | 0.00 | 0.00 | 0.00 |
| CV**IET** | 0.00 | 0.41 | 0.66 | 0.14 | 0.93 | 1.00 | 0.75 | 0.00 |
| CV**IDT** | 0.00 | 0.00 | 0.00 | 0.00 | 0.00 | 0.00 | 0.16 | 0.00 |
| **Y**V**IET** | 0.00 | 0.00 | 0.00 | 0.00 | 0.00 | 0.00 | 0.00 | 0.00 |
| *Other* | 0.00 | 0.00 | 0.00 | 0.00 | 0.00 | 0.00 | 0.05 | 0.00 |
| All with **T** | 1.00 | 0.41 | 0.66 | 0.14 | 0.93 | 1.00 | 0.97 | 0.99 |

**Supplementary Table 12. Frequencies of *dhfr* (51, 59, 108, 164) and *dhps* (437, 540, 581, 613) multi-locus haplotypes.** Mutant amino acids are shown in **underlined bold** font, wild-type in normal font. The first row shows the wild-type NCSI/AKAA haplotype. The following rows shows 61 distinct mutant homozygous haplotypes, ordered by the number of mutations. The proportions in these rows are proportions amongst all samples that had a homozygous haplotype. In addition to these haplotypes, many samples had heterozygous haplotypes, or the full haplotypes could not be resolved. These are shown in the Other row as a proportion of all samples. Note that many haplotypes are rare with only IRNI/AKAA, IRNI/GKAA, IRNI/GEAA, IRNL/GEAA, IRNL/GEGA and IRNL/GNGA having frequency > 5%. SAM=South America, WAF=West Africa, CAF=Central Africa, EAF=East Africa, SAS=South Asia, WSEA=West south-east Asia, ESEA=East south-east Asia, OCE=Oceania.

| DHFR/DHPS haplotype | SAM (n=37) | WAF (n=1565) | CAF (n=260) | EAF (n=635) | SAS (n=50) | WSEA (n=963) | ESEA (n=1077) | OCE (n=195) | All (n=4782) |
|---|---|---|---|---|---|---|---|---|---|
| NCSI/AKAA | 0.03 | 0.07 | 0.00 | 0.01 | 0.00 | 0.00 | 0.00 | 0.01 | 0.03 |
| NCSI/**G**KAA | 0.00 | 0.08 | 0.00 | 0.00 | 0.00 | 0.00 | 0.00 | 0.00 | 0.03 |
| NC**N**I/AKAA | 0.59 | 0.00 | 0.00 | 0.00 | 0.00 | 0.00 | 0.00 | 0.00 | 0.00 |
| NCSI/AKA**S** | 0.00 | 0.00 | 0.00 | 0.00 | 0.00 | 0.00 | 0.00 | 0.00 | 0.00 |
| NC**T**I/AKAA | 0.00 | 0.00 | 0.00 | 0.00 | 0.00 | 0.00 | 0.00 | 0.00 | 0.00 |
| N**RN**I/AKAA | 0.00 | 0.01 | 0.00 | 0.00 | 0.02 | 0.00 | 0.03 | 0.38 | 0.03 |
| **I**C**N**I/AKAA | 0.08 | 0.00 | 0.00 | 0.01 | 0.00 | 0.00 | 0.00 | 0.00 | 0.00 |
| NCSI/**G**KA**S** | 0.00 | 0.01 | 0.00 | 0.00 | 0.00 | 0.00 | 0.00 | 0.00 | 0.00 |
| NC**N**I/**G**KAA | 0.03 | 0.00 | 0.00 | 0.00 | 0.00 | 0.00 | 0.00 | 0.00 | 0.00 |
| NCSI/**GE**AA | 0.00 | 0.00 | 0.00 | 0.00 | 0.00 | 0.00 | 0.00 | 0.00 | 0.00 |
| **IRN**I/AKAA | 0.00 | 0.17 | 0.04 | 0.03 | 0.00 | 0.00 | 0.09 | 0.00 | 0.08 |
| N**RN**I/**G**KAA | 0.00 | 0.04 | 0.00 | 0.00 | 0.00 | 0.00 | 0.03 | 0.05 | 0.02 |
| **I**C**N**I/**G**KAA | 0.03 | 0.01 | 0.15 | 0.00 | 0.00 | 0.00 | 0.00 | 0.00 | 0.01 |
| NC**N**I/**G**K**G**A | 0.05 | 0.00 | 0.00 | 0.00 | 0.00 | 0.00 | 0.00 | 0.00 | 0.00 |
| **I**C**N**I/AKA**S** | 0.00 | 0.00 | 0.00 | 0.00 | 0.00 | 0.00 | 0.00 | 0.00 | 0.00 |
| NC**N**I/**GE**AA | 0.00 | 0.00 | 0.00 | 0.00 | 0.00 | 0.00 | 0.00 | 0.01 | 0.00 |
| NC**N**I/**G**KA**S** | 0.00 | 0.00 | 0.00 | 0.00 | 0.00 | 0.00 | 0.00 | 0.00 | 0.00 |
| N**RNL**/AKAA | 0.00 | 0.00 | 0.00 | 0.00 | 0.02 | 0.00 | 0.00 | 0.00 | 0.00 |
| **IRN**I/**G**KAA | 0.00 | 0.53 | 0.73 | 0.02 | 0.04 | 0.00 | 0.16 | 0.00 | 0.25 |
| N**RN**I/**GE**AA | 0.00 | 0.00 | 0.00 | 0.02 | 0.30 | 0.02 | 0.01 | 0.55 | 0.03 |
| **I**C**N**I/**GE**AA | 0.00 | 0.00 | 0.02 | 0.03 | 0.02 | 0.00 | 0.00 | 0.00 | 0.01 |
| **IRN**I/AKA**S** | 0.00 | 0.01 | 0.00 | 0.00 | 0.00 | 0.00 | 0.00 | 0.00 | 0.00 |
| N**RN**I/**G**KA**S** | 0.00 | 0.00 | 0.00 | 0.00 | 0.00 | 0.00 | 0.00 | 0.00 | 0.00 |
| **IRNL**/AKAA | 0.00 | 0.00 | 0.00 | 0.00 | 0.00 | 0.00 | 0.01 | 0.00 | 0.00 |
| N**RN**I/**G**K**G**A | 0.00 | 0.00 | 0.00 | 0.00 | 0.00 | 0.00 | 0.00 | 0.00 | 0.00 |
| N**RNL**/**G**KAA | 0.00 | 0.00 | 0.00 | 0.00 | 0.00 | 0.00 | 0.00 | 0.00 | 0.00 |
| **I**C**N**I/**G**KA**S** | 0.00 | 0.00 | 0.00 | 0.00 | 0.00 | 0.00 | 0.00 | 0.00 | 0.00 |
| N**RN**I/**G**KA**T** | 0.00 | 0.00 | 0.00 | 0.00 | 0.00 | 0.00 | 0.00 | 0.00 | 0.00 |
| **IRN**I/**GE**AA | 0.00 | 0.01 | 0.03 | 0.75 | 0.16 | 0.04 | 0.17 | 0.00 | 0.15 |
| **IRN**I/**G**KA**S** | 0.00 | 0.03 | 0.00 | 0.00 | 0.00 | 0.00 | 0.00 | 0.00 | 0.01 |
| N**RNL**/**GE**AA | 0.00 | 0.00 | 0.00 | 0.00 | 0.06 | 0.03 | 0.00 | 0.00 | 0.01 |

| | | | | | | | | | |
|---|---|---|---|---|---|---|---|---|---|
| NRNI/GEGA | 0.00 | 0.00 | 0.00 | 0.00 | 0.04 | 0.02 | 0.00 | 0.00 | 0.00 |
| IRNL/GKAA | 0.00 | 0.00 | 0.00 | 0.00 | 0.00 | 0.00 | 0.02 | 0.00 | 0.00 |
| IRNI/GKGA | 0.00 | 0.00 | 0.00 | 0.01 | 0.00 | 0.00 | 0.01 | 0.00 | 0.00 |
| ICNI/GEGA | 0.08 | 0.00 | 0.01 | 0.00 | 0.00 | 0.00 | 0.00 | 0.00 | 0.00 |
| NRNI/GNGA | 0.00 | 0.00 | 0.00 | 0.00 | 0.00 | 0.00 | 0.00 | 0.00 | 0.00 |
| NRNL/GNAA | 0.00 | 0.00 | 0.00 | 0.00 | 0.00 | 0.00 | 0.00 | 0.00 | 0.00 |
| ICNL/GKGA | 0.08 | 0.00 | 0.00 | 0.00 | 0.00 | 0.00 | 0.00 | 0.00 | 0.00 |
| NRNI/GKGS | 0.00 | 0.00 | 0.00 | 0.00 | 0.00 | 0.00 | 0.00 | 0.00 | 0.00 |
| NRNL/GKGA | 0.00 | 0.00 | 0.00 | 0.00 | 0.00 | 0.00 | 0.00 | 0.00 | 0.00 |
| IRNI/GKAT | 0.00 | 0.00 | 0.00 | 0.00 | 0.00 | 0.00 | 0.00 | 0.00 | 0.00 |
| ICNL/GEAA | 0.00 | 0.00 | 0.00 | 0.00 | 0.00 | 0.00 | 0.00 | 0.00 | 0.00 |
| IRNI/AKGS | 0.00 | 0.00 | 0.00 | 0.00 | 0.00 | 0.00 | 0.00 | 0.00 | 0.00 |
| IRNI/GNAA | 0.00 | 0.00 | 0.00 | 0.00 | 0.00 | 0.00 | 0.00 | 0.00 | 0.00 |
| IRNI/GYAA | 0.00 | 0.00 | 0.00 | 0.00 | 0.00 | 0.00 | 0.00 | 0.00 | 0.00 |
| NRNL/GIAA | 0.00 | 0.00 | 0.00 | 0.00 | 0.00 | 0.00 | 0.00 | 0.00 | 0.00 |
| IRNL/GEAA | 0.00 | 0.00 | 0.00 | 0.00 | 0.18 | 0.13 | 0.09 | 0.00 | 0.05 |
| IRNI/GEGA | 0.00 | 0.00 | 0.02 | 0.11 | 0.02 | 0.06 | 0.02 | 0.00 | 0.03 |
| IRNI/GNGA | 0.00 | 0.00 | 0.00 | 0.00 | 0.00 | 0.01 | 0.07 | 0.00 | 0.02 |
| IRNI/GKGS | 0.00 | 0.02 | 0.00 | 0.00 | 0.00 | 0.00 | 0.00 | 0.00 | 0.01 |
| NRNL/GEGA | 0.00 | 0.00 | 0.00 | 0.00 | 0.04 | 0.02 | 0.00 | 0.00 | 0.00 |
| IRNL/GKGA | 0.00 | 0.00 | 0.00 | 0.00 | 0.04 | 0.01 | 0.01 | 0.00 | 0.00 |
| IRNI/GEAS | 0.00 | 0.00 | 0.00 | 0.00 | 0.00 | 0.00 | 0.01 | 0.00 | 0.00 |
| IRNL/GNAA | 0.00 | 0.00 | 0.00 | 0.00 | 0.00 | 0.00 | 0.00 | 0.00 | 0.00 |
| NRNL/GEAT | 0.00 | 0.00 | 0.00 | 0.00 | 0.00 | 0.00 | 0.00 | 0.00 | 0.00 |
| IRNI/GEAT | 0.00 | 0.00 | 0.00 | 0.00 | 0.00 | 0.00 | 0.00 | 0.00 | 0.00 |
| ICNL/GEGA | 0.03 | 0.00 | 0.00 | 0.00 | 0.00 | 0.00 | 0.00 | 0.00 | 0.00 |
| IRNL/GYAA | 0.00 | 0.00 | 0.00 | 0.00 | 0.00 | 0.00 | 0.00 | 0.00 | 0.00 |
| IRNL/GEGA | 0.00 | 0.00 | 0.00 | 0.00 | 0.06 | 0.58 | 0.01 | 0.00 | 0.12 |
| IRNL/GNGA | 0.00 | 0.00 | 0.00 | 0.00 | 0.00 | 0.05 | 0.22 | 0.00 | 0.06 |
| IRNL/GEAS | 0.00 | 0.00 | 0.00 | 0.00 | 0.00 | 0.00 | 0.03 | 0.00 | 0.01 |
| IRNL/GEAT | 0.00 | 0.00 | 0.00 | 0.00 | 0.00 | 0.00 | 0.00 | 0.00 | 0.00 |

| | | | | | | | | | |
|---|---|---|---|---|---|---|---|---|---|
| **Other** | 0.00 | 0.30 | 0.24 | 0.14 | 0.35 | 0.11 | 0.15 | 0.03 | 0.20 |
| **(n/total)** | (0/37) | (666/2231) | (84/344) | (104/739) | (27/77) | (116/1079) | (185/1262) | (6/201) | (1188/5970) |

**Supplementary Table 13. Frequency of HRP2 and HRP3 deletions by country.** n=number of samples for which an unambiguous HRP deletion genotype (deleted or non-deleted) could be assigned.

| Country | HRP2 deletions | HRP3 deletions | HRP2 and HRP3 deletions |
|---|---|---|---|
| Bangladesh (n=77) | 0% | 0% | 0% |
| Benin (n=36) | 0% | 0% | 0% |
| Burkina Faso (n=56) | 0% | 0% | 0% |
| Cambodia (n=896) | 0% | 3% | 0% |
| Cameroon (n=235) | 0% | 0% | 0% |
| Colombia (n=16) | 0% | 0% | 0% |
| Congo DR (n=344) | 0% | 0% | 0% |
| Ethiopia (n=21) | 0% | 43% | 0% |
| Gambia (n=219) | 0% | 0% | 0% |
| Ghana (n=851) | 0% | 0% | 0% |
| Guinea (n=149) | 0% | 0% | 0% |
| Indonesia (n=80) | 4% | 25% | 0% |
| Ivory Coast (n=70) | 0% | 0% | 0% |
| Kenya (n=110) | 0% | 1% | 0% |
| Laos (n=120) | 0% | 1% | 0% |
| Madagascar (n=24) | 0% | 0% | 0% |
| Malawi (n=254) | 0% | 0% | 0% |
| Mali (n=426) | 0% | 0% | 0% |
| Mauritania (n=76) | 0% | 0% | 0% |
| Mozambique (n=1) | 0% | 0% | 0% |
| Myanmar (n=211) | 0% | 0% | 0% |
| Nigeria (n=29) | 0% | 0% | 0% |
| Papua New Guinea (n=121) | 0% | 0% | 0% |
| Peru (n=21) | 38% | 67% | 29% |
| Senegal (n=84) | 0% | 7% | 0% |
| Tanzania (n=316) | 0% | 0% | 0% |
| Thailand (n=888) | 0% | 0% | 0% |
| Uganda (n=13) | 0% | 0% | 0% |
| Viet Nam (n=226) | 0% | 4% | 0% |

**Supplementary Table 14. Alleles at six mitochondrial positions used for the species identification.** The loci are all located within the *cox3* gene. Nucleotide pairs in square brackets indicate that either allele at that position is a match.

| **Locus** | **Positions** | **Allele by Species** | | | | | |
|---|---|---|---|---|---|---|---|
| | | ***P. falciparum*** | ***P. vivax*** | ***P. knowlesi*** | ***P. malariae*** | ***P. ovale wallikeri*** | ***P. ovale curtisi*** |
| 1 | 668-671 | ATGA | TTTA | TTTT | TTGT | ATTT | ATTT |
| | 678-683 | TTGT[CT]T | TATTAT | TATTAT | ATTAAT | ACATAA | ATATAT |
| 2 | 728-733 | GTTCAT | TATTCA | | GTTCAA | GTTACA | |
| | 740-740 | T | T | | T | A | |
| | 749-751 | TAA | AAA | | TAG | TAA | |
| | 770-773 | GA[TC]T | TACA | | TACT | TATT | |
| 3 | 861-869 | TCGGTAGAA | TCACTATTA | TCACAATTA | TCACTATTT | CCCTTATTT | TCGTTATTA |
| | 878-881 | TATT | CATT | AACT | AATA | AACT | AACT |
| | 884-887 | TATT | AACT | TATT | TATC | AACC | AACC |
| 4 | 971-982 | AGTATATACAGT | ACCAGATATAGC | ACCTGATATAGC | TCCTGAAACTCC | ACCAGATATAGC | |
| 5 | 1025-1028 | TAGA | AAGT | | TAAT | TAAT | TAAT |
| | 1046-1049 | TAAT | AAGT | | AAGT | AAGA | AAGG |
| 6 | 1062-1066 | CAAAT | AATA[CT] | | AATAT | AATAT | |
| | 1073-1073 | A | A | | T | T | |
| | 1076-1076 | G | A | | T | T | |
| | 1082-1082 | A | T | | A | T | |
| | 1091-1091 | T | T | | A | T | |
| | 1102-1108 | TAAATAC | TTAGAAA | | AAAGAAA | T[GA]AGAAA | |

### Supplementary figures

**Supplementary Figure 1. Histogram of local differentiation score for all genes**. Red line shows the 99th percentile. A selection of known drug resistance genes are marked, all of which have high local differentiation score. fd=PF3D7_1318100 (putative ferredoxin).

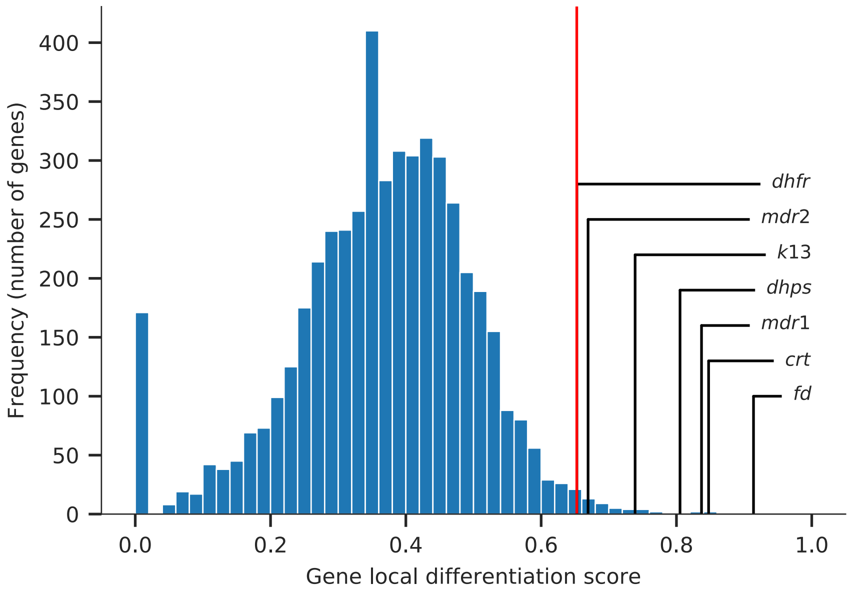
